# Reference frames for encoding of translation and tilt in the caudal cerebellar vermis

**DOI:** 10.1101/2023.09.09.556993

**Authors:** Félix Buron, Christophe Z. Martin, Jessica X. Brooks, Andrea M. Green

## Abstract

Many daily behaviors rely critically on estimates of our body’s motion and orientation in space. Vestibular signals are essential for such estimates but to contribute appropriately two key sets of computations are required. First, ambiguous motion information from the otolith organs must be combined with spatially transformed rotational signals (e.g., from the canals) to distinguish head translation from tilt. Second, tilt and translation estimates must be transformed from a head- to a body-centered reference frame to correctly interpret the body’s motion. Studies have shown that cells in the caudal cerebellar vermis (nodulus and ventral uvula, NU) reflect the output of the first set of computations to estimate translation and tilt. However, it remains unknown whether these estimates are encoded exclusively in head-centered coordinates or whether they reflect a further transformation towards body-centered coordinates. Here we addressed this question by examining how the 3D spatial tuning of otolith and canal signals on translation- and tilt-selective NU neurons varies with changes in head-re-body and body-re-gravity orientation. We show that NU cell tuning properties are consistent with head-centered coding of otolith signals during translation. Furthermore, while canals signals in the NU have been transformed into world-referenced estimates of reorientation relative to gravity (i.e., tilt), as needed to resolve the tilt-translation ambiguity, the resulting tilt estimates are encoded in head-centered coordinates. Our results thus suggest that body-centered motion and orientation estimates required for postural control, navigation and reaching are computed elsewhere either by further transforming NU outputs or via computations in other parallel pathways.

## Introduction

Daily tasks such as navigating, controlling posture and planning voluntary movements require knowledge of the motion and orientation of the body. Vestibular signals are essential for estimating body motion, but to contribute appropriately at least two key problems must be resolved. First, because our otolith organs transduce inertial (translational) and gravitational (tilt) accelerations equivalently (Fernandez and Goldberg, 1976a,b; Angelaki and Dickman, 2000), distinguishing translation from tilt requires combining otolith signals with rotational signals that provide an independent estimate of reorientation relative to gravity (Merfeld et al., 1999; Green and Angelaki, 2004, 2007; Laurens and Angelaki, 2017). Second, because the vestibular sensors are fixed in the head, the way they are activated by a given body motion depends on the head’s movement and orientation with respect to the body. Vestibular and neck proprioceptive signals must thus be integrated to correctly estimate body motion (Kleine et al., 2004; Shaikh et al., 2004; Brooks and Cullen, 2009; Luan et al., 2013; Martin et al., 2018; Zobeiri and Cullen, 2022).

An interconnected brainstem-cerebellar network has been implicated in resolving both problems (Green and Angelaki, 2010). Neurons in the vestibular (VN) and rostral fastigial (rFN) deep cerebellar nuclei (Angelaki et al., 2004; Green et al., 2005; Shaikh et al., 2005b; Mackrous et al., 2019) as well as upstream thalamic (Meng et al., 2007) and cortical areas (Liu et al., 2011) reflect distributed or population-level estimates of translation. Furthermore, a solution to the otolith “tilt/translation ambiguity” is reflected in individual neurons of the caudal cerebellar vermis (nodulus and ventral uvula, NU, lobules X, IXc,d) where distinct translation- and tilt-coding Purkinje cells have been identified (Yakusheva et al., 2007; Laurens et al., 2013; Stay et al., 2019; Hernández et al., 2020). In line with theoretical predictions (e.g., Green and Angelaki, 2004,2007), NU neurons combine otolith signals with canal signals that have been spatially transformed into estimates of the earth-horizontal component of rotation that indicates head reorientation relative to gravity (i.e., tilt; Yakusheva et al., 2007; Laurens et al., 2013). However, to be useful for tasks such as maintaining balance or voluntary reaching, vestibular-derived estimates of head tilt and translation must ultimately be integrated with proprioceptive cues to estimate body motion and orientation.

Compelling evidence suggests that the rFN plays an important role in encoding such body motion estimates. Many rFN neurons integrate vestibular and proprioceptive signals to distinguish head from body motion (Brooks and Cullen, 2009, 2013) and reflect the required transformation of head-centered vestibular signals towards body-centered coordinates (Kleine et al., 2004; Shaikh et al., 2004; Martin et al., 2018). Recently, Martin et al. (2018) examined this transformation for the first time fully in three-dimensions (3D) to show that body-centered representations in 3D could be linearly decoded from small populations of rFN cells. On the basis of this and other observations it was suggested that the rFN may reflect a late stage in the computations, with the bulk of the nonlinear computations required to effect the transformation occurring upstream in the cerebellar cortex. This notion is supported by previous findings implicating regions of the anterior vermis (AV) that project to the rFN in transforming vestibular signals towards body-centered coordinates (Manzoni et al., 1998; Manzoni et al., 1999). Yet, most motion-sensitive rFN cells also reflect at least a partial solution to the tilt/translation ambiguity (Angelaki et al., 2004; Green et al., 2005; Shaikh et al., 2005b; Martin et al., 2018; Mackrous et al., 2019) and thus the output of computations thought to be performed largely upstream in the NU of the posterior vermis. Consequently, if both sets of computations occur upstream of the rFN in the cerebellar cortex a key question arises: Do NU cells encode a solution to the tilt/translation ambiguity only in head-centered coordinates or are tilt and translation estimates in this region further transformed towards body-centered coordinates? Here we addressed this question by examining for the first time how the spatial tuning for translation and tilt in the NU varies with changes in head-re-body orientation.

## Methods

### Animal preparation

Data reported here were collected from 3 male rhesus monkeys (Macaca mulatta, 7-12 kg) that were prepared for chronic recording of eye movements and single-unit activity. In a first surgery, animals were chronically implanted with a 7-cm diameter delrin head-stabilization ring that was anchored to the skull using hydroxyapatite-coated inverted T-bolts and neuro-surgical acrylic bone cement (Martin et al., 2018). Supports for a removable delrin recording grid (3×5 cm) were stereotaxically positioned inside the ring and secured to the skull with acrylic. The recording grid consisted of arrays of holes spaced by 0.8 mm arranged in staggered rows. Holes on each side of the grid were slanted relative to the horizontal plane 10° upward from lateral to medial to provide bilateral access to medial cerebellar regions. In a second surgery, animals were chronically implanted with a scleral search coil for recording eye movements in 2D (Robinson, 1963). After surgical recovery, animals were trained to fixate and pursue visual targets within a ±1.5° window for fluid reward using standard operant conditioning techniques. All surgical procedures were performed aseptically and under anesthesia. Surgical and experimental procedures were approved by the institutional animal research review board (Comité de déontologie de l’expérimentation sur les animaux) and were in accordance with national guidelines for animal care.

### Experimental setup

During experiments, the animals were comfortably seated in custom-built primate chairs that were designed to permit the head to be secured in different positions relative to the body by means of a movable head restraint system (Martin et al., 2018). This head restraint system, which attached to the animal’s ring implant at three points via setscrews, allowed the head to be manually reoriented along three independent orthogonal axes and locked in different head-on-trunk positions (Fig. 1A). In particular, the head could be reoriented in vertical planes toward nose-down (pitch reorientation by up to 45°; Fig. 1Aii,Dii) and right/left-ear-down (roll reorientation by up to 30°; Fig. 1Aiii,Diii) about horizontal axes located at approximately the level of the second cervical vertebra. The head could also be reoriented in the horizontal plane about a vertical axis passing through the head center (yaw axis reorientation by up to 45°; Fig. 1Aiv, Div). To prevent changes in torso and limb position the animal’s torso was secured with shoulder straps and a waist restraint and his limbs were gently secured to the chair using soft forearm and ankle sleeves.

**Figure 1.**
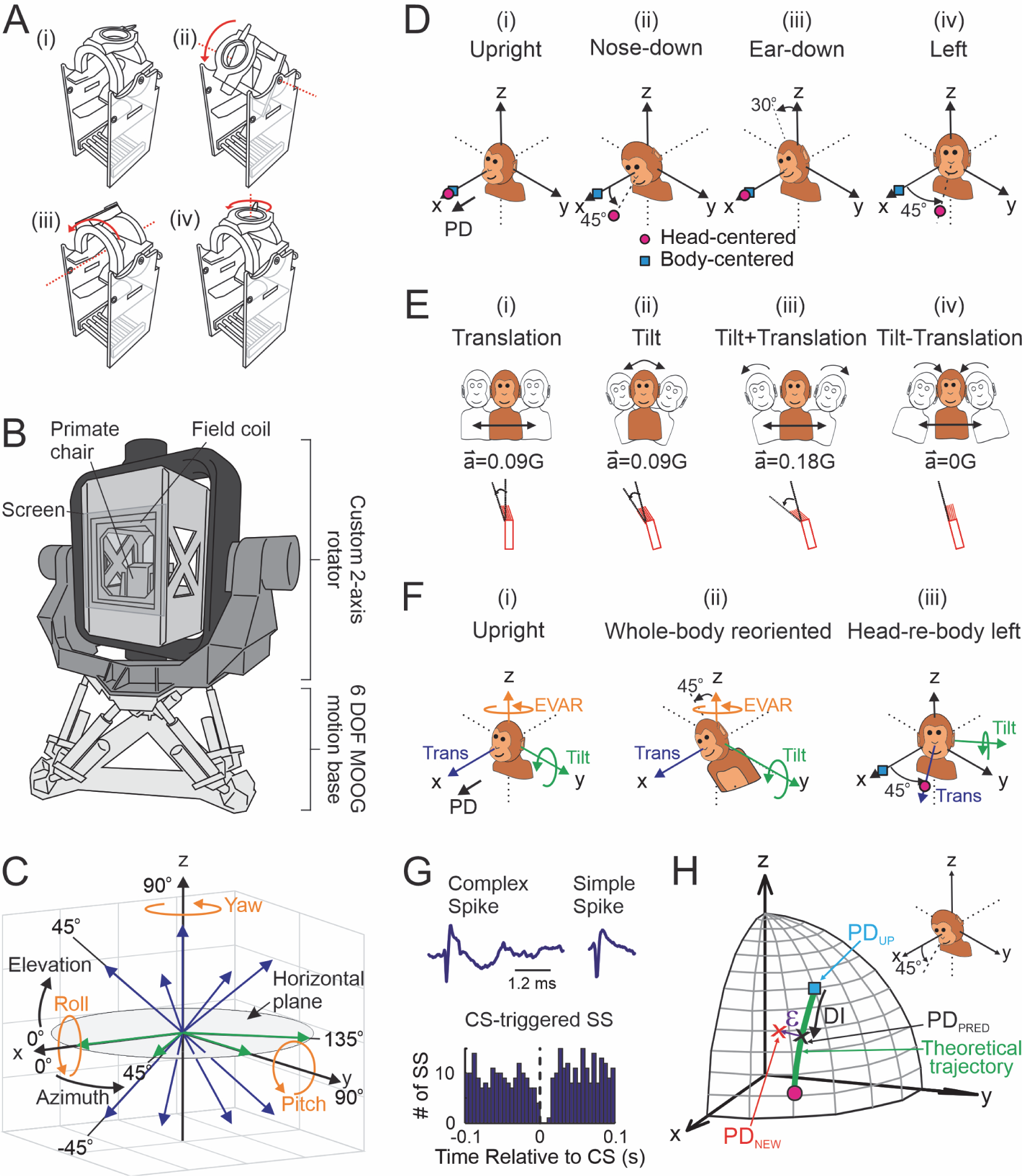
Experimental methods. **A**, Custom primate chairs used to test cell tuning with the head upright (**Ai**) as well as after manual repositioning of the head relative to the body in the vertical plane by up to 45° toward nose-down (**Aii**) or 30° toward right-ear-down (**Aiii**) and in the horizontal plane by up to 45° to the left (**Aiv**). **B**, Schematic illustration of the motion delivery system consisting of a custom 2-axis rotator (Actidyn) mounted on top of a 6 degree-of-freedom motion platform (Moog). The primate chair and eye monitoring system (field coil) were mounted to the innermost axis of the rotator system. **C**, Illustration of the 13 axes along which sinusoidal translation stimuli (blue and green arrows; 0.5Hz ±9cm, ±0.09G) and the 3 axes about which sinusoidal rotation stimuli (orange arrows; 0.5Hz, ±31.4°/s) were applied to characterize cell response tuning. **D**, Cell tuning for translation was characterized with the head upright (**Di**), after vertical plane head reorientation relative to the body by either 45° toward nose-down (**Dii**) or 30° toward ear-down (**Diii**) and/or after horizontal plane reorientation 45° to the left (**Div**). The predicted preferred direction (PD) for body-centered (blue squares) versus head-centered (pink circles) encoding of translation is illustrated for a cell with a head upright PD along the x axis (0° azimuth, 0° elevation). **E**, Tilt and translation stimulus combinations used to test a subset of cells along the 4 horizontal plane axes (green arrows in **C**). These included sinusoidal tilts (**Eii**; 0.5Hz, ±5.19°, ±0.09G) matched in amplitude to the translation stimulus (**Ei**; 0.5Hz ±9cm, ±0.09G) as well simultaneous translation and tilt stimuli combined in either an additive (**Eiii**; Tilt + Translation, 0.5 Hz, ±0.18G) or subtractive (**Eiv**; Tilt - Translation, 0.5 Hz, ±0G) fashion. **F**, Cells were tested during earth-vertical-axis rotations (EVAR; orange) and for the stimuli in **E** applied along the horizontal-plane axis closest to the cell’s PD for the Translation-Tilt stimulus when upright (**Fi**), as well as after the whole body was reoriented relative to gravity by 45° along the axis perpendicular to the cell’s Translation-Tilt PD (e.g., after body reorientation towards right/left-ear-down for testing along the 0° azimuth PD axis; **Fii**) and after head reorientation by 45° to the left (**Fiii**). **G**, NU Purkinje cells were identified by the presence of both simple spikes (SS) and complex spikes (CS; top) and by showing that simple spike activity paused for at least 10 ms after the occurrence of a complex spike (bottom). **H**, Schematic representation of the theoretically predicted trajectory of PDs (green curve) between body-centered (blue square) and head-centered (pink circle) coding predicted for an example cell with an upright PD (PD_UP_) of (45°, 45°). The cell’s tuning reflects a new PD (PD_NEW_; red x) of (25°, 30°) when the head is reoriented 45° toward nose-down (inset). The extent of shift from PD_UP_ to PD_NEW_ along the “ideal” (green) trajectory was quantified by a displacement index (DI; x corresponds to a DI of 0.5). The extent of shift orthogonal to this trajectory was quantified by an angular error ε (purple arrows illustrate an ε of −8.5°) defined as the angle between PD_NEW_ (red x) and the closest point along the ideal trajectory (PD_PRED_; black x).

The primate chair was secured to a motion system that was used to provide 3D translational and rotational motion stimuli. In initial experiments, motion stimuli were delivered using a 6 degree-of-freedom motion platform described previously (6DOF2000E, Moog; see Martin et al., 2018). This system was subsequently extended to incorporate a custom-built gimballed 2-axis rotator system (Actidyn Inc.) that was integrated with and mounted on top of the motion platform (Fig. 1B). The gimballed rotator system consists of an inner “yaw-axis” rotator mounted inside an outer “pitch-axis” rotator. The primate chair was secured to the innermost frame of the rotator such the animal’s body-vertical axis was aligned with the inner yaw rotation axis and the center of the animal’s head (intersection of his interaural and naso-occipital head axes) was positioned at the intersection of the two rotator axes (i.e., center of the gimballed rotator system). Consequently, rotation of the innermost axis of the rotator reoriented the animal about his body-vertical axis while rotation about the outer pitch axis reoriented his body in a vertical plane with respect to gravity. Importantly, the complete 3D motion system allows delivery of different combinations of 3D translational and rotational motion stimuli across a broad range of different body orientations with respect to gravity.

The motion stimuli delivered to the animal were measured using a navigational sensor composed of a three-axis linear accelerometer and a three-axis angular velocity sensor (Tri-Axial Navigational IMU, Kistler) which was mounted to the chair support inside the innermost frame, just adjacent to the animal’s head. Eye movements were measured with a three-field magnetic search coil system (24-inch cube; Riverbend Instruments) that was mounted on the innermost frame of the rotator such that the monkey’s head was centered within the magnetic field. Visual targets were back-projected onto a vertical screen mounted in front of the animal (30 cm distance) using a wall-mounted laser and x-y mirror galvanometer system (Cambridge Technology). Stimulus presentation, reward delivery, and data acquisition were controlled with custom scripts written in the Spike2 software environment using the Cambridge Electronics Design (model power 1401) data acquisition system. Eye coil voltage signals and 3D linear accelerometer and angular velocity measurements of the motion stimuli delivered to the animal were antialias filtered (200 Hz, 4-pole Bessel; Krohn-Hite), digitized at a rate of 833.33 Hz, and stored using the Cambridge Electronics Design system.

### Neural recording

Single-unit extracellular recordings in the caudal cerebellar vermis (lobules IX and X; nodulus and ventral uvula) were performed with epoxy-coated, etched tungsten microelectrodes (3-5MΩ, FHC) that were inserted into the brain using 26-gauge guide tubes that were passed through the recording grid and small predrilled burr holes in the skull. A remote-controlled mechanical microdrive (Hydraulic Probe Drive; FHC) mounted on the animal’s head-stabilization ring was then used to advance electrodes. Neural activity was amplified, filtered (30 Hz to 15 kHz), and isolated online using a time-amplitude dual window discriminator (BAK Electronics). Single-unit spikes triggered acceptance pulses from the window discriminator that were time-stamped and stored using the event channel of the Cambridge Electronics Design data acquisition system. In addition, raw neural activity waveforms were digitized at 30 kHz and stored using the Cambridge Electronics Design system for off-line spike discrimination.

In initial penetrations the abducens nuclei and rostral fastigial nuclei were localized bilaterally. The regions of interest in the nodulus and ventral uvula (NU) were then identified on the basis of their stereotaxic location in relation to the abducens nuclei, fastigial nuclei and fourth ventricle (e.g., Yakusheva et al., 2007). Motion-sensitive NU cells were recorded within ± 4.3 mm of the midline (92% within 2.5mm), 4.8 mm to 12 mm posterior to the abducens nuclei and at depths of 3.5 mm above to 2.8 mm below the abducens. Recordings were restricted to neurons in the Purkinje cell layer where both simple and complex spikes could be observed and heard on the audio monitor. Among recorded cells, 74% (70/95) were explicitly identified offline as Purkinje cells by the capacity to identify the characteristic waveforms of both simple spikes and complex spikes and by showing that simple spike activity paused for at least 10 ms after the occurrence of a complex spike (Fig. 1G; Barmack and Shojaku, 1995). Units for which both simple and complex spikes were present in the recording, but clear simple spike pausing was not observed were considered putative Purkinje cells. Because we found no differences between identified and putative Purkinje cell responses to motion stimuli across the conditions tested (see below) the two groups have been presented together in the results. All cells described here were characterized as eye-movement-insensitive by their failure to exhibit changes in modulation during horizontal and vertical smooth pursuit (0.5 Hz, ±10 cm) as well as during saccades and fixation (up to ±20° horizontally and vertically). Cell responses to motion stimuli were recorded in darkness and eye position was uncontrolled.

After a putative NU Purkinje cell was isolated, its responses to motion stimuli were first characterised with the animal’s head facing straight ahead and with the head and body in an upright orientation. The cell’s 3D spatial tuning for translation was characterised by recording responses to sinusoidal translation (0.5Hz, ±9cm, ±0.09G) along 13 different axes. These axes were defined in body/world-centered coordinates and described in a spherical coordinate system in terms of combinations of azimuth and elevation angles in increments of 45° spanning 3D space (Fig. 1C; blue and green arrows). The cell’s 3D rotation tuning was also characterised during sinusoidal rotation (0.5Hz, ±31 deg/s, ±10 deg) about the x, y and z axes (Fig. 1C, orange arrows). The majority of neurons (N=69/95) were further characterized along the 4 horizontal-plane axes (azimuth 0°, 45°, 90°, 135° for elevation 0°; Fig. 1C, green arrows) for different combinations of matched translation and tilt stimuli. These stimuli have been used in previous studies (Fig. 1E; Angelaki et al., 2004; Green et al., 2005; Yakusheva et al., 2007, Mackrous et al., 2019; Hernández et al., 2020) to evaluate how central neurons integrate otolith and canal signals to distinguish components of the net gravito-inertial acceleration (GIA), sensed by the otoliths, due to translational motion from those due to head reorientation relative to gravity. In addition to the purely translational motion stimuli described above, these stimuli consisted of pure tilts matched to the translation stimulus (see below; Fig. 1Eii) as well as simultaneous translational and tilt motion stimuli combined in either an additive (Tilt+Translation; Fig. 1Eiii) or subtractive (Tilt-Translation; Fig. 1Eiv) fashion. In particular, pure tilts consisted of 0.5 Hz sinusoidal rotations from upright with a peak amplitude of ±5.19° (±16.3 deg/s), producing a peak GIA stimulus to the otoliths that was matched in magnitude (0.09 G) to that produced by the pure translation. During combined tilt and translation stimuli inertial and gravitational components of linear acceleration combined along a given axis either additively to produce double the net GIA (Tilt+Translation stimulus; 0.18 G) or subtractively such that the net GIA was close to zero (Tilt-Translation stimulus; 0 G). Importantly, because the Tilt-Translation stimulus produces little activation of otolith afferents (i.e, net GIA close to 0; Angelaki et al., 2004) it provides a means of isolating semicircular canal contributions to central neural activities during earth-horizontal-axis rotations that dynamically reorient or tilt the head with respect to gravity.

To examine the reference frames in which translation-sensitive NU cells encode otolith signals, spatial tuning for translation was subsequently characterized in one or more additional head-re-body positions obtained by statically reorienting the head in vertical and/or horizontal planes (Fig. 1D). In particular, head reorientation in the vertical plane consisted of either pitch towards 45° nose-down or roll towards 30° right-ear-down, depending on the cell’s preferred translation direction when upright and the axis of reorientation predicted to result in the larger differences between head- and body-centered tuning (i.e., the axis further away from the cell’s preferred upright translation direction; Fig. 1Dii vs. 1Diii). Reorientation of the head in the horizontal plane involved displacement of the head to the left by 45° (Fig. 1Div). After statically positioning the head in a new orientation, neural responses were characterized for the same set of body/world-centered translation directions that were examined with the head upright and facing forward. Because tuning shifts that can distinguish between reference frames are predicted to be largest in the plane of head reorientation, translation directions in this plane were always tested first to maximize the potential to use partial data sets. After characterizing cell response tuning in each new head orientation, the head was returned to the upright and straight forward position and responses along one or more directions close to the cell’s preferred response direction were re-tested to confirm cell isolation before further characterization of the cell’s activity.

To examine the reference frames in which canal-derived estimates of tilt are encoded within the NU, additional tests were performed on a subset of recorded neurons whose responses were characterized across different body-re-gravity and/or head-re-body orientations for the matched tilt and combined tilt/translation stimuli described above. In particular, we sought to 1) explicitly confirm that the canal signals carried by these cells had been spatially transformed into a true 3D estimate of tilt relative to gravity; and 2) investigate the reference frames in which such tilt signals are encoded. Cells carrying true canal-derived tilt estimates should respond similarly to the Tilt-Translation stimulus (i.e., which isolates canal signals during earth-horizontal-axis rotation) across different whole-body orientations relative to gravity. Consequently, to address the first issue we reoriented the whole body of the animal relative to gravity by 45° along the horizontal plane axis (i.e., azimuth 0°, 45°, 90° or 135°) closest to perpendicular to its preferred axis for the Tilt-Translation stimulus (e.g., towards right-ear-down for a PD closest to 0°; Fig. 1Fi,ii). The cell’s responses to earth-vertical-axis rotation (EVAR) as well as to translation, tilt, Tilt-Translation and Tilt+Translation along the preferred Tilt-Translation direction were then retested. Three cells were also retested for whole-body reorientation along the same axis but in the opposite direction (e.g., towards left-ear-down for a PD of 0°). Of particular relevance is the fact that across all body orientations (i.e., upright and body tilted) translation and tilt stimuli were applied along the same horizontal-plane axis in head/body coordinates (e.g., 0° azimuth in Fig. 1Fi). Consequently, the dynamic stimulus to the otoliths was similar in both whole-body orientations. In contrast, stimuli involving rotation activated different sets of canals in each body orientation. For example, tilts stimulated mainly the vertical canals when upright but both horizontal and vertical canals after body reorientation. Conversely, EVAR stimulated mainly the horizontal canals when upright and both vertical and horizontal canals after body reorientation. Importantly, cells that had been spatially transformed into a world-referenced estimate of dynamic tilt should not reflect these different sensory activation patterns. Instead, they should exhibit similar responses to the Tilt-Translation stimulus and remain unresponsive to EVAR across the different whole-body orientations.

Finally, to address the second issue (i.e., reference frames in which canal-derived estimates of tilt are encoded) head and body reference frames were dissociated by reorienting the head relative to the body by 45° to the left in the body upright orientation (Fig. 1Fiii). Responses to the translation, tilt and combined tilt and translation stimuli were then retested.

### Data analysis

Data analysis was performed offline using custom scripts written in MATLAB (The MathWorks). Spike times were converted into instantaneous firing rate (IFR) by taking the inverse of the interspike interval and assigning this value to the middle of the interval. The gain and phase of the neural IFR response to each motion stimulus in a given direction and head/body orientation were obtained by fitting 5-20 well-isolated cycles of both the response and the stimulus with a sine function using a nonlinear least-squares minimization procedure (Levenberg-Marquardt). Confidence intervals (CIs) on estimates of neural response gain and phase were obtained using a bootstrap method in which bootstrapped gain and phase estimates were obtained by resampling (with replacement) the response cycles used for the sine function fits 600 times. A cell was considered to exhibit a significant modulation for a given motion stimulus if at the time of peak response (estimated from the non-bootstrapped fit across response cycles) 95% CIs on the firing rate were greater than a minimum of 2 spikes/sinusoidal cycle. For translation stimuli, response gain was expressed in units of spikes/s/G (where G = 9.81 m/s^2^) and phase as the difference (in degrees) between peak neural modulation and peak translational acceleration. For rotational stimuli, gain was expressed in units of spikes/s/deg/s and phase as the difference between peak neural modulation and peak angular velocity. To facilitate comparison of cell responses to translation with those due to tilt relative to gravity, earth-horizontal-axis rotation responses were also calculated relative to the linear acceleration stimulus to the otoliths caused by tilt in units of spikes/s/G with phase expressed as the difference between peak neural modulation and peak acceleration.

Cells were subsequently classified based on their horizontal-plane response gains and phases as predominantly encoding the net GIA (i.e., like the otoliths), translation, tilt or having composite properties (i.e., not better correlated with either translation or tilt). Using a multiple linear regression analysis, neural responses to translation, tilt and combined tilt/translation stimuli (whenever possible) across directions in the horizontal plane were first fit with a general model which assumes that cell responses can be described as a weighted combination of acceleration components due to translation and tilt. Specifically, we fit the model *Response* = H_1_*Trans*_x_ + H_2_*Trans*_y_ + H_3_*Tilt*_x_ + H_4_*Tilt*_y_ where *Response* is a vector of the observed response amplitudes and phases (expressed compactly as a complex number) for the different stimuli, and *Trans_x_*, *Trans_y_*, *Tilt_x_*, and *Tilt_y_* are the corresponding accelerations due to translation and tilt associated with each stimulus along the x (azimuth 0°) and y (azimuth 90°) axes. Regression coefficients, H_1_-H_4_, are complex numbers embedding both gain and phase information which provide estimates of the cell’s responsiveness to translation and tilt along each axis. In addition to the general model, we also fit two simpler models including a “translation model” which assumes that the neuron responds selectively to translation (i.e., H_3_, H_4_ =0) and a “tilt model” which assumes that the neuron responds selectively to tilt (i.e., H_1_, H_2_ =0). Each model was fit 600 times using the bootstrapped response gains and phases for each stimulus, described above, to obtain distributions of model parameters (H1-H4). These were in turn used to compute distributions of tuning functions describing cell spatio-temporal sensitivities to translation and/or tilt (Angelaki, 1991).

To identify cells whose responses were consistent with encoding the net GIA we compared the tuning functions for translation versus tilt that were obtained from fits using the general model. Specifically, we calculated distributions of differences in tuning function parameters (gains, phases and preferred direction) for translation versus tilt. Cells whose 95% CIs (percentile method) for gain, phase and preferred direction differences included zero were classified as GIA-encoding. For comparison, we also calculated distributions of gain, phase and preferred direction differences for translation versus tilt obtained by fitting responses to pure translation and tilt stimuli only using the simpler translation and tilt models and obtained the same results.

Cells not classified as GIA-encoding were then further examined to determine whether they could be considered statistically better correlated with either translation or tilt. We compared the fits of first-order translation and tilt models by calculating the correlation between the data and each set of model predictions as well as the correlation between the model predictions themselves. These were used to compute partial correlation coefficients which were then converted to normalized partial-correlation scores (Z-scores) and used to assess significant differences in the model fits (see *Statistical Analysis* below). Cells that were significantly better correlated with the translation than the tilt model were classified as translation-coding. Cells were classified as tilt-encoding if they were both significantly better correlated with the tilt model and had responses to vertical-axis translation that were either negligible or smaller than those for horizontal-plane translations. All other cells were classified as composite.

The spatial tuning of each cell was subsequently characterized across one or more head orientations. The tuning profiles of cells classified as being responsive to translation (i.e., all cells except those identified as preferentially encoding tilt) were visualized in 3D by plotting response gains as a function of the azimuth and elevation of translation direction (e.g., Fig. 3A-C). A precise estimate of the cell’s preferred response direction (PD) for translation was obtained by fitting response gains and phases across directions using a 3D spatio-temporal convergence (STC) model (e.g., Fig. 3D; for details see Chen-Huang and Peterson, 2006; Martin et al., 2018) that is more general than cosine tuning and can account for the observation that many brainstem and cerebellar neurons exhibit translation responses that vary in terms of both gain and phase with motion direction (Siebold et al., 1999; Angelaki and Dickman, 2000; Shaikh et al., 2005a; Chen-Huang and Peterson, 2006; Martin et al., 2018). These fits yielded estimates of the cell’s maximum response gain, phase and direction (i.e., PD). In addition, we calculated the ratio of minimum to maximum response gain (STC tuning ratio) which provides a measure of the extent to which a cell’s response reflects simple cosine-tuned changes in gain across motion directions (STC ratio = 0) versus dynamic properties that vary with motion direction (STC ratio > 0), consistent with the convergence of vestibular signals with different preferred response directions and phases relative to the stimulus (Angelaki, 1991; Chen-Huang and Peterson, 2006). The majority (96%) of cells responsive to translation (i.e., cells other than tilt-selective cells) were fit with a 3D STC model with the head upright. In addition, 84% of datasets collected after either horizontal or vertical plane head reorientation were fit with this model. For the remaining datasets insufficient data was collected after head reorientation to fit the full 3D model. Such partial datasets were included only if we were able to complete testing across 4 directions in the plane of head reorientation. In these cases, preferred tuning was estimated by fitting responses across motion directions in that plane using a 2D STC model (Angelaki, 1991). In addition, since tilt, by definition, implies rotation about an earth-horizontal-plane axis, the spatial tuning for tilt was characterized in the horizontal plane using a 2D STC model. We defined the tilt PD as the horizontal-plane direction along which body tilt produced the maximum response.

To examine the extent to which cells encoded motion in a head- or body-centered reference frame, we quantified how their spatial tuning varied across changes in head-re-body orientation. Because movement directions were defined in body/world-centered coordinates, cells encoding motion in a body-centered reference frame should exhibit spatial tuning that remains invariant to changes in head-re-body orientation (e.g., Fig 1D, blue squares). In contrast, head-centered cells should shift their spatial tuning with changes in head position to an extent that could be predicted based on the cell’s PD and the amplitude and plane of head reorientation. For cells whose tuning properties were characterised fully in 3D across multiple head orientations, we quantified such spatial tuning shifts using a 3D displacement index (DI). This index, defined as the ratio of the actual change in spatial tuning compared with that predicted for head-centered tuning (ΔPD_actual_/ΔPD_head-centered_), simultaneously took into account both the azimuth and elevation components of observed and predicted shifts (i.e., full 3D shifts in tuning) for head reorientation in a given plane (see Martin et al., 2018 for details). No shift in PD with changes in head orientation thus yielded a DI of 0 (body-centered tuning) whereas a PD shift consistent with the predictions for head-centered tuning yielded a DI of 1. In brief, the DI was obtained by first calculating an “ideal trajectory” of predicted PDs that would be obtained if the neuron’s tuning shifted by different fractions of the way towards the full shift predicted for a head-centered cell (Fig 1H; green trace). The cell’s DI for head reorientation in a given plane was then obtained by computing the dot product between the actual PD observed after head reorientation (Fig. 1H; PD_NEW,_ red cross) and each of the predicted PDs along the trajectory to find which predicted PD (corresponding to a particular DI) was the closest match to the observed PD (Fig. 1H; PD_PRED,_ black cross). Because a given PD shift could also have components along a direction orthogonal to the “ideal trajectory” we also estimated the angular deviation of the observed PD away from the closest predicted PD. This was done by calculating an angular error, ε, between these two directions as 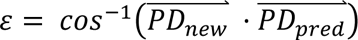 with clockwise versus counterclockwise angular deviation from the ideal PD trajectory between head and body-centered coordinates defined as positive and negative errors, respectively. For cells in which the translational spatial tuning after a given head reorientation was characterized only in 2D, the DI was calculated based on the ratio of observed to predicted shifts in the plane of head reorientation only (i.e., no ε values were calculated). Similarly, spatial tuning shifts for tilt were characterized in 2D based on the ratio of observed to predicted shifts in the horizontal plane.

To establish the significance of tuning shifts we computed confidence intervals (CIs) for each DI (and ε values, when applicable) by using a bootstrap method based on resampling of residuals (Efron and Tibshirani, 1993). In brief, bootstrapped tuning functions for each head orientation were obtained by resampling (with replacement) the residuals of the STC model fit to the data, adding the resampled residuals to the model predicted response to create a new synthetic dataset and then fitting this dataset with the STC model to obtain a new set of model parameters. This procedure was repeated 1000 times to obtain a distribution of tuning functions and corresponding distributions of DIs (and ε values) in each head orientation from which 95% CIs were obtained (percentile method). A DI (or ε) was considered significantly different from a particular value if its 95% CI did not include that value.

The DI analysis yielded an estimate of the extent of transformation toward body-centered coordinates for head reorientations in a given plane. In addition, to determine whether a cell could be considered more consistent with head- versus body-centered encoding of translation we performed a second model-fitting analysis. Specifically, we simultaneously fit data collected across multiple head orientations with two different 3D STC models. These included a body-centered reference frame model in which response gains and phases were expressed in terms of motion directions defined in body-centered coordinates, and a head-centered model in which gains and phases were instead expressed in head-centered coordinates. The fitting procedure was similar to that used to quantify 3D spatial tuning in a single head orientation (see above) but when extended across multiple head orientations incorporated an additional gain scaling factor to take into account any global changes in cell response gain (i.e., a general gain increase/decrease across all motion directions) associated with different head orientations (see Martin et al., 2018, for details). We then estimated the goodness of fit of each model by computing the correlation (R) between the model prediction and the data. Because the two models are themselves correlated, we removed the influence of this correlation by computing partial correlation coefficients (ρ) (see Equations 1 and 2 in *Statistical Analysis* below) which were then normalized using Fisher’s r-to-Z transform. Score differences equivalent to p < 0.05 (see below) were considered significant. To visualize the results of this analysis, we constructed a scatterplot of Z-scores for the head-centered versus body-centered model fits and separated the plot into regions in which one model fit was significantly better than that of the other (e.g., see Fig. 5).

### Statistical analysis

All statistical tests were performed in MATLAB (The MathWorks). Bootstrap methods were used to evaluate the significance of cell modulation in response to a given stimulus as well as to evaluate the significance of tuning shifts. In brief, to establish CIs on the estimates of neural response gain and phase obtained from sine function fits to the firing rate modulation for a given motion stimulus and direction (see above), we used a bootstrap method in which bootstrapped gain and phase estimates were obtained by resampling (with replacement) the response cycles used for the sine function fits 600 times. A cell was considered to exhibit a significant modulation for a given motion stimulus if at the time of peak response 95% CIs on the firing rate modulation were greater than a minimum of 2 spikes/sinusoidal cycle. The significance of changes in response tuning (i.e., estimated using STC models) was evaluated by using a bootstrap method based on resampling of residuals (Efron and Tibshirani, 1993) to create distributions of 1000 tuning functions for each head orientation. These were used to compute corresponding distributions of PD response gain and phase changes, STC ratio changes, DIs, and *ℇ* values from which 95% CIs could be derived (for details, see above; Martin et al., 2018). A given parameter was considered significantly different from a particular value if the 95% CI did not include that value.

To test whether neural responses were better described by a “translation” versus “tilt” model as well as whether cell responses across head orientations were better correlated with a “head-centered” versus “body-centered” tuning model we used partial correlation analyses which took into account correlations between the competing models themselves. In particular, to distinguish between competing models we first computed the correlation between each model and the data. We then removed the influence of correlations between the two competing models by computing partial correlation coefficients according to the equations:

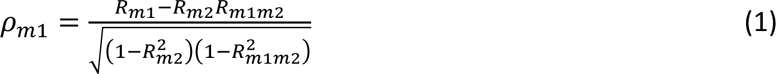

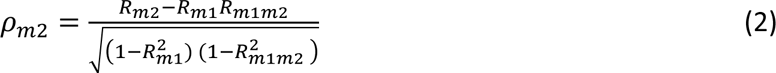

where *R_m1_* and *R_m2_*, are the correlation coefficients between the data and each of competing models 1 and 2 (i.e., “translation” vs “tilt” or “head-centered” vs “body-centered”) and *R_m1m2_* is the correlation between the two models. Partial correlation coefficients *ρ_m1_* and *ρ_m2_* were then normalized using Fisher’s r-to-Z transform to allow conclusions regarding significance to be made on based on comparison of Z-scores, independent of the number of data points (Angelaki et al., 2004; Smith et al., 2005; Martin et al., 2018). A cell’s response was considered significantly better fit by one model as compared to the other model if the Z score for that model was greater than 1.645 and exceeded the Z score of the other model by 1.645 (equivalent to p<0.05).

We tested the uniformity of translational PDs (i.e., circular distributions of azimuth and elevation angles) in our sampled neural populations using Rao’s spacing test (Berens, 2009). The Wilcoxon rank-sum test was used to test for significant differences between DI distributions. The Wilcoxon signed rank test was used to compare distribution medians to specific values (e.g., DI medians to either 0 or 1). Hartingan’s Dip Test (Hartigan and Hartigan, 1985) was used to test distributions for multimodality. Correlations between DI and other cell tuning properties (Fig. 6) and between response gains across body orientations (Fig. 9A) were evaluated using linear regression. For all statistical tests p-values of less than 0.05 were considered statistically significant.

## Results

We recorded the responses of 95 confirmed or putative Purkinje cells in the nodulus/uvula (NU) of the caudal vermis (42 in Monkey A, 27 in Monkey B, 26 in monkey C) that were insensitive to eye movements but responsive to passive movement stimuli provided by a 3D motion system (Fig. 1B). To characterize their simple spike responses, we recorded neural activity as monkeys were sinusoidally translated along 13 directions spanning 3D space (Fig. 1C; blue and green lines). Most cells (86/95) were also characterized during rotations about the three cardinal (x,y,z) axes (Fig. 1C, orange arrows) and 73% (69/95) were further characterized during combined tilt/translation stimuli (“Tilt+Translation” and “Tilt-Translation” stimuli; see Methods) in 4 horizontal-plane directions (Fig. 1C; green arrows). Based on these responses, cells were classified either as responding similarly to tilt and translation (i.e., like otolith afferents) such that they encoded net gravito-inertial acceleration (GIA), or as preferentially encoding either translation or tilt. Cells that were neither GIA-encoding, nor better fit by either translation or tilt models were classified as “composite”. Comparison of the horizontal-plane sensitivities of our recorded cells to tilt versus translation (Fig. 2A) revealed that a majority of classified cells exhibited substantially larger responses to translation as compared to tilt with 68% of cells (59/87) being classified as preferentially translation-encoding (Fig. 2A, filled blue circles; mean tilt/translation gain ratio of 0.27) while 22% (19/87) were classified as tilt-encoding (Fig. 2A, filled green circles; mean tilt/translation gain ratio of 4.2). Notably, only two cells were classified based on their horizontal-plane responses as GIA-encoding (Fig. 2A, orange circles) and both had substantially larger sensitivities to vertical-axis translation than to horizontal-plane tilt and translation stimuli. All remaining cells were characterized as composite (8%, 7/87; Fig. 2A, filled gray circles). Thus, in agreement with previous studies (Yakusheva et al., 2007; Laurens et al., 2013; Laurens and Angelaki, 2020) most Purkinje cells in the NU reflected a solution to the tilt/translation ambiguity such that they preferentially encoded either translation or tilt.

**Figure 2.**
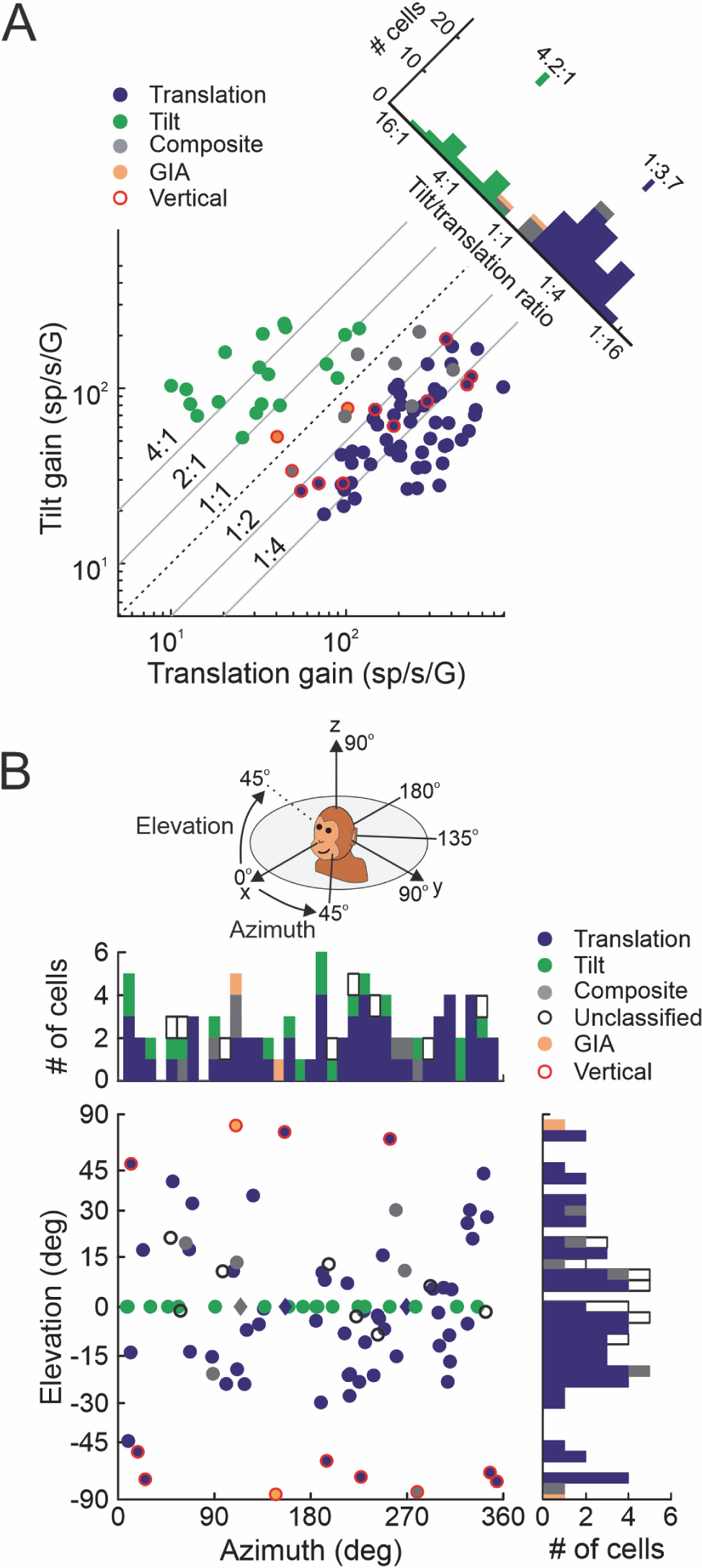
Cell selectivity for tilt versus translation and distribution of preferred response directions. **A**, Comparison of cell horizontal-plane response gains for translation versus tilt. Cells are color-coded according to classification type as preferentially translation-coding (blue filled; N=59), tilt-coding (green, N=19), GIA-coding (orange; N=2) or composite (gray; N=7). Cells with responses to translation along the vertical (0°, 90°) axis that were larger than their horizontal-plane responses to both tilt and translation were identified as “vertical” translation cells and outlined in red (N=12). Plotted gains for translation and tilt correspond to estimates in the cell’s maximum sensitivity direction for each stimulus in the horizontal plane. The histogram at top right shows the distribution of tilt/translation gain ratios. Bars above the plot indicate mean values. **B**, Distribution of preferred directions for translation for all translation-sensitive NU neurons whose tuning was characterized in 3D (i.e., including translation-coding, composite and unclassified cells; N=73). Each data point in the scatter plot indicates the preferred azimuth (abscissa) and elevation (ordinate) with elevation angle spacing scaled according to Lambert’s equal area projection of the spherical stimulus space. Histograms along the top and right sides of the plot show marginal distributions. Black open symbols denote cells that could not be classified. All other colors correspond to the classifications in **A**. The preferred horizontal-plane (azimuth) tilt directions for tilt-selective neurons (green circles along 0° elevation; N=19) and translation-sensitive neurons whose tuning was only characterized in 2D (diamonds along 0° elevation; N=3) are also shown.

The spatial tuning properties of all translation-sensitive neurons (i.e., all neurons except those classified as tilt-encoding) were visualized by plotting their responses as a function of translation direction in azimuth and elevation to construct 3D tuning functions (e.g., Fig. 3A). Precise estimates of their preferred direction (PD) for translation were computed by fitting neural responses with a 3D STC model (see Materials and Methods; Fig. 3D). As illustrated in Figure 2B, translation PDs (Fig. 2B, filled blue circles) spanned 3D space when the head was upright and facing forward. PDs were uniformly distributed across azimuth directions but were largely concentrated in elevation within ±45° of the horizontal plane (Rao’s spacing test, azimuth p=0.5, elevation p=0.001). Likewise, the tilt PDs of all tilt-sensitive neurons (Fig. 2B, filled green circles), estimated using a 2D STC model, were uniformly distributed in the horizontal plane (Rao’s spacing test, p=0.5).

**Figure 3.**
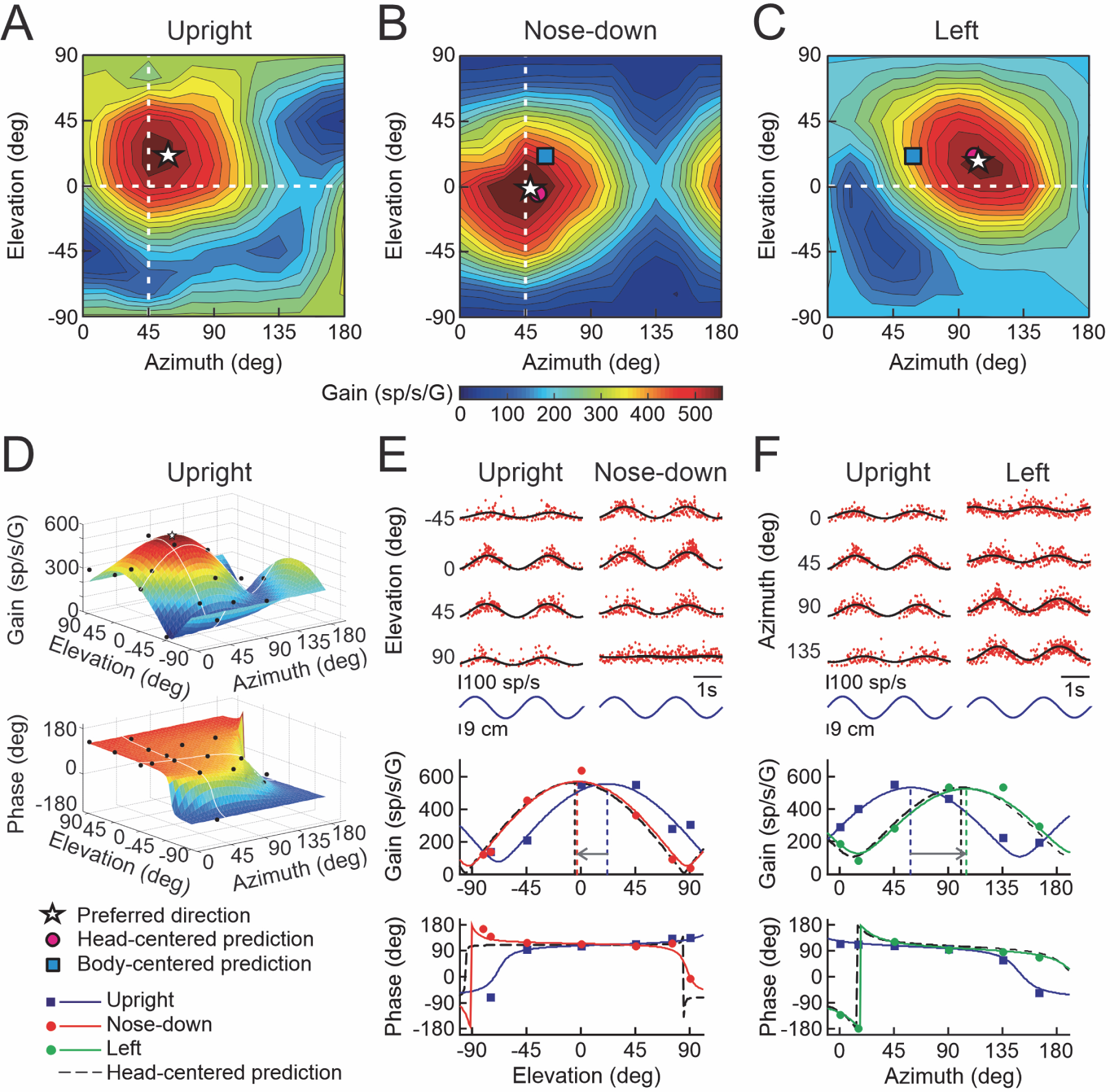
Example NU Purkinje cell with head-centered tuning for translation. **A-C**, Contour plots illustrating the cell’s 3D response tuning for translation with the head upright (**A**) as well as after vertical-plane head reorientation 45° toward nose-down (**B**) and after horizontal-plane reorientation 42° to the left (**C**). Azimuth and elevation are expressed in body/world-centered coordinates. Star: PD; pink circle: predicted PD for head-centered tuning; blue square: predicted PD for body-centered tuning. **D**, The cell’s preferred translation direction (star) for each head orientation was estimated by fitting a 3D STC model to cell response gain and phase data (black circles) across stimulus directions (here illustrated for the upright orientation). **E**, Instantaneous firing rates (IFRs; top) and STC model fit to gains and phases across elevations (bottom) for an azimuth angle of 45° (i.e., along dashed white line in **B**) with the head upright (blue lines and symbols) and after reorientation toward nose-down (red lines and symbols). **F**, IFRs (top) and STC model fit to gains and phases across azimuth angles (bottom) for 0° elevation (i.e., along dashed white line in **C**) with the head upright (blue lines and symbols) and after reorientation to the left (green lines and symbols). Dashed black curves in **E** and **F** show the predictions for head-centered tuning.

### Reference frames for encoding otolith signals during translation

To investigate the reference frames in which NU Purkinje cells encode otolith signals during translation, head- and body-centered reference frames were dissociated by comparing spatial tuning with the head upright to that after head reorientation relative to the body in vertical and/or horizontal planes (Fig. 1D). Because translation directions were defined in a coordinate frame that remained fixed to the body/world, the spatial tuning of a neuron encoding translation in body-centered coordinates should be invariant to changes in head-re-body position (Fig. 1D, blue squares). In contrast, cells encoding translation in head-centered coordinates (i.e., like vestibular afferents) should exhibit systematic shifts in preferred tuning with changes in head orientation (Fig. 1D, pink circles).

To quantify spatial tuning shifts across the neural population we computed displacement indices (DIs) for each cell and plane of head reorientation that expressed the extent of the cell’s tuning shift as a fraction of that predicted for head-centered tuning. In the majority of cases (N=41/49 head reorientations) PD shifts were quantified fully in 3D, taking into account the extent of shift in both azimuth and elevation along the “ideal trajectory” for a transformation between head and body-centered coordinates (i.e., the DI; see Fig. 1H, green trace). In addition, we quantified the angular displacement orthogonal to this trajectory (Fig. 1H, angular deviation error, ε; see Materials and Methods). A DI of 1 (and ε of 0) indicates a tuning shift equivalent to that predicted for head-centered tuning, whereas a DI of 0 indicates no shift, consistent with body/world-centered coding.

Figure 3 illustrates a representative NU cell. With the head in the upright orientation, the preferred direction of the cell was 58.8° in azimuth and 20.9° in elevation (Fig. 3A, white star). However, after vertical-plane head-re-body reorientation by 45° towards nose-down, the cell’s PD shifted by −23.5° in elevation and −10.7° in azimuth. This is close to the predictions for head-centered encoding of translation as illustrated in the contour plot of Figure 3B (star closer to pink circle than blue square) and STC model fits to the data across elevations in the 45° azimuth plane (Fig. 3E, bottom; red curve close to back dashed curve). Similarly, after head reorientation in the horizontal plane by 42° to the left, the cell’s PD shifted by 45.1° in the same direction as the head (Fig. 3C,F). Shifts in both planes were consistent with encoding of translation in a mainly head-centered reference frame as reflected by DIs close to 1 (DI_vert_=0.90; DI_hor_=1.07) and small ε values (ε_vert_=5°; ε_hor_=4°).

Similar to the example cell of Figure 3, NU cells generally exhibited tuning properties that were more consistent with head-centered coding. This is shown in Figure 4 which summarizes the distributions of DIs (Fig. 4A) and ε values (Fig. 4C, green) for all NU cells tested across head reorientations in the vertical and/or horizontal planes. Also shown for comparison are the DI and ε distributions for rFN and VN cells from the prior study of Martin et al. (2018) (Fig. 4B and C, bottom; red: rFN; blue: VN). DI distributions for NU cells reflected median DI values that were close to 1 for reorientation in both planes (Fig. 4A; DI_vert_ = 0.79, DI_hor_ = 0.94) while mean absolute ε values were close to 0 (Fig. 4C, top: mean ε = 4.7° across both planes). In line with this, 87% (20/23) of DI values for horizontal-plane reorientation were statistically indistinguishable from 1 and while this percentage was slightly lower for vertical plane reorientation (62%, 16/26) there was no significant difference in the DI distributions for reorientation in vertical versus horizontal planes (Wilcoxon rank-sum test, p = 0.11). Notably, DI values for NU cells closely resembled those previously reported for rostral VN cells which were found to be predominantly head-centered (Martin et al., 2018; Fig. 4B, blue). No significant difference was found in the DI distributions of the two populations (Wilcoxon rank-sum test, p = 0.94). In contrast, the DI distribution of rFN cells was substantially more broadly distributed between body- and head-centered reference frames (Fig. 4B, red) with a median DI value (0.61) that was significantly lower than that of NU cells (Wilcoxon rank-sum test, p < 0.001). NU cell translation responses thus reflected a coding of otolith signals that was significantly more head-centered than that of rFN neurons.

**Figure 4.**
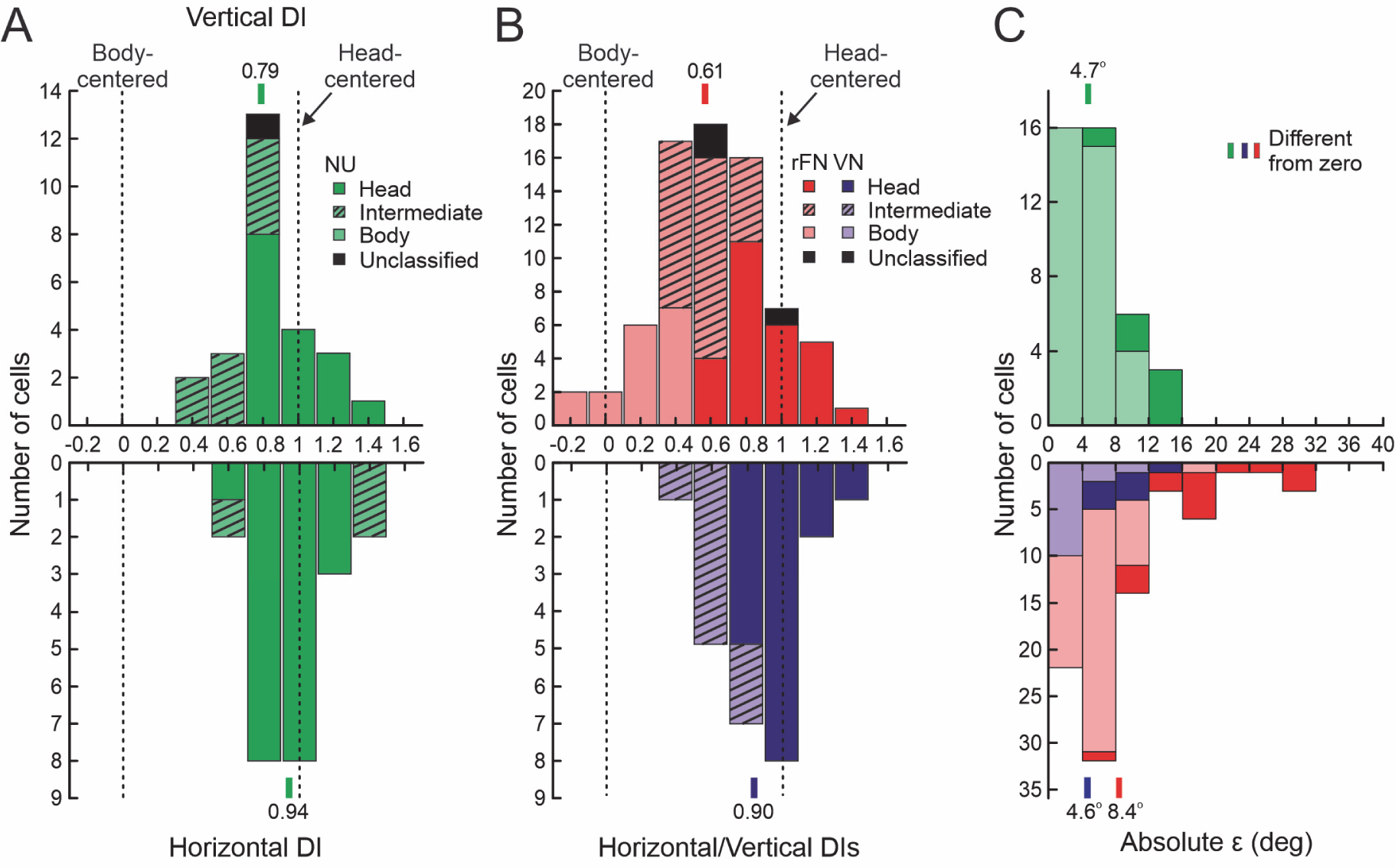
Distribution of DI and ε values for the spatial tuning of otolith signals during translation across changes in head-re-body orientation. DI values of 0 and 1 indicate tuning shifts consistent with body- and head-centered encoding of otolith signals, respectively. **A**, DI distribution for NU cells for vertical-plane (top, N=26) and horizontal-plane (bottom, N=23) head reorientations. **B**, Combined vertical- and horizontal-plane DI distributions for rFN cells (red; vertical, N=47; horizontal, N=27 horizontal) and VN cells (blue; vertical, N=16; horizontal, N=8) replotted from the study of Martin et al. (2018). DIs in **A** and **B** are classified based on bootstrapped DI 95% CIs as head-centered (dark colors), body-centered (pale colors) or intermediate (hatched). Cells for which bootstrapped DIs included both 0 and 1 were labelled as unclassified (black). Vertical bars above each plot indicate the medians of the populations. **C**, Distributions of absolute ε values across both vertical- and horizontal-plane head orientations for NU cells (green, top) compared with those of rFN (red) and VN (blue) cells (bottom) from the study of Martin et al. (2018). Dark colors indicate a significant difference from zero (N=6 of 41 cases for NU; N=16 of 62 cases for rFN; N=7 of 20 cases for VN). Vertical bars above the plot indicate the means of each distribution.

This distinction in tuning properties across brainstem-cerebellar regions was further emphasized when responses were compared using a model-fitting analysis. Specifically, the DI analysis provided a model-independent characterization of tuning shifts. Confidence intervals on DI values obtained through bootstrap analyses (see Materials and Methods) provided an indication of whether those shifts could be considered statistically different from either head- or body-centered coding (i.e., DI of 1 or 0). However, a limitation of this approach for classifying cells is that the size of the confidence intervals on our DI values was influenced by the quality of our STC fits in each head orientation. Consequently, a cell with a DI close to but slightly less than 1 (e.g., 0.8) could be classified as intermediate (i.e., DI statistically different from both 1 and 0) if the STC function provided an excellent fit (and very narrow DI confidence intervals) while a second cell with the same DI but more variable responses and a poorer STC function fit would be more likely to be classified as head-centered (i.e., DI not statistically different from 1). Furthermore, because the DI analysis considered tuning shifts in only one plane of head reorientation at a time it thus provided little insight as to how a given cell should be classified when considered fully in 3D. Thus, to address these issues we performed a second model-fitting analysis that provided another means of classifying cell spatial tuning properties. This analysis assessed whether the tuning of each cell was best explained by a body- versus a head-centered model by fitting neural responses with each of these models across multiple head orientations (see Materials and Methods).

For comparison with the DI results, we first fit each model to the data across two head orientations in a single plane (i.e., across upright and ear/nose-down orientations or across upright and left orientations). The goodness of fit of each model was quantified using a partial correlation analysis. To facilitate interpretation and visualization of the results, partial correlation coefficients (normalized using Fisher’s r-to-Z transform) for fits of the head-centered model were plotted versus those for the body-centered model (Fig. 5A). Cells in the upper-left region of the plot were significantly better fit by the head-centered model, whereas those in the lower-right region were significantly better fit by a body-centered model. Cells falling in the central diagonal region were not better fit by one model as compared to the other and were classified as “intermediate”.

**Figure 5.**
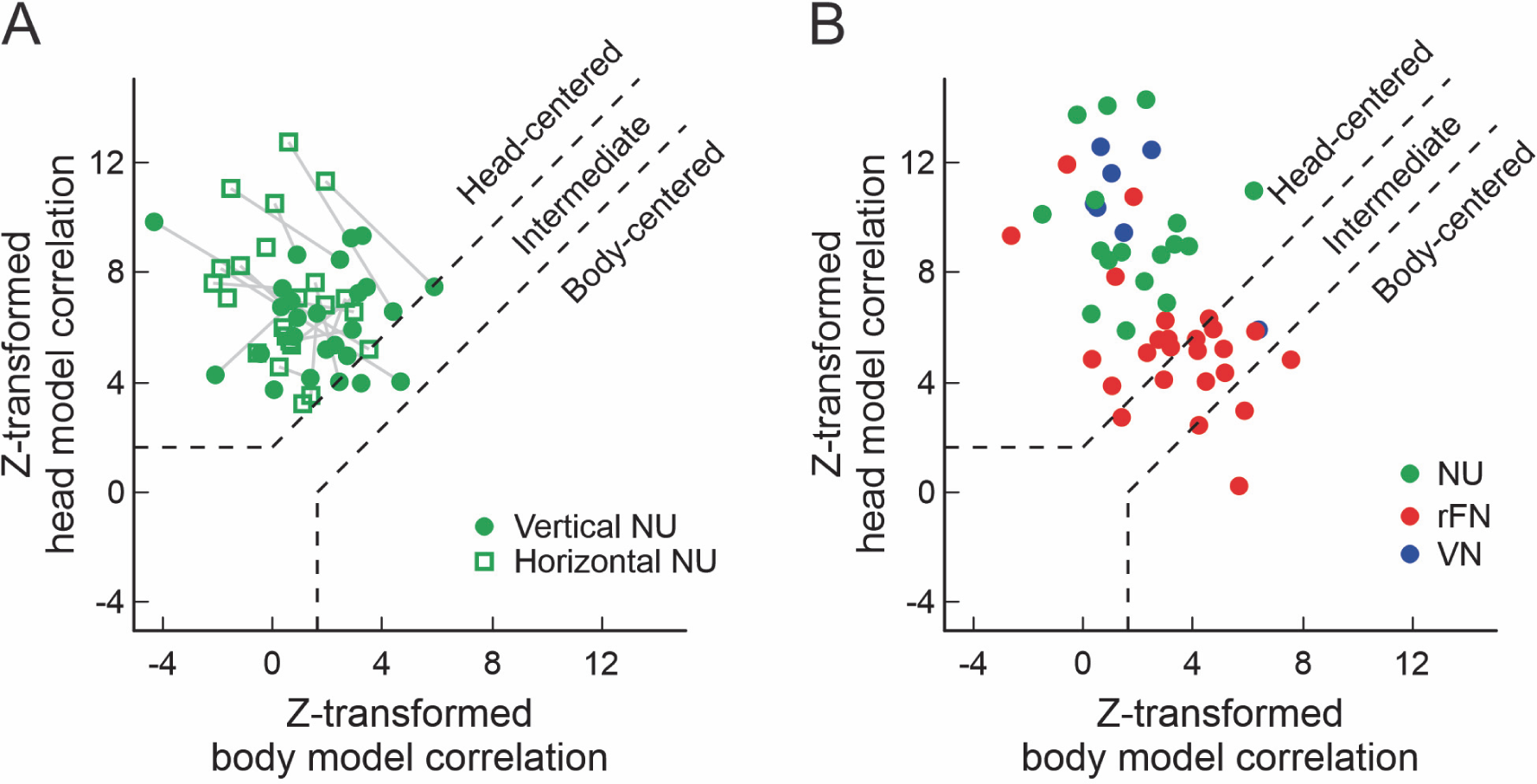
Correlation of otolith spatial tuning with head-centered and body-centered models. **A**, **B**, Scatter plots of head- versus body-centered model partial correlation coefficients (Z-transformed) when translation response gains and phases for NU cells (green) were fit with each model across either (**A**) 2 head orientations (circles: upright and after vertical plane reorientation, N=26; squares: upright and after horizontal plane reorientation, N=23) or (**B**) 3 head orientations (upright and after vertical and horizontal plane reorientations, N=17). Corresponding results for model fits to rFN (red, N=25) and VN (blue, N=7) cells from the study of Martin et al. (2018) are shown superimposed in **B** for comparison. Significance regions are based on the difference between head- and body-centered model fit Z-scores corresponding to p<0.05 (top left: head-centered; bottom right: body-centered; middle diagonal region: not significantly better correlated with either model and classified as “intermediate”). Gray lines in **A** link data points associated with the same cell for head reorientation in different planes.

Using this approach, we found that all (100%; 23/23) NU neurons were better fit by a head-centered model for horizontal plane head reorientation (Fig. 5A, open green squares). Similarly, for vertical-plane reorientation a majority of cells (85%; 22/26) were classified as head-centered while the remaining 15% (4/26) were classified as intermediate (Fig. 5A, filled green circles). The predominance of head-centered translation coding was further emphasized when the model-fitting analysis was extended across all 3 head orientations for the subset of cells (N=17) characterized for reorientations in both vertical and horizontal planes. The results of this analysis are shown in Figure 5B, superimposed on the results of a similar analysis previously reported for rFN and rostral VN cells (Martin et al., 2018). All NU cells characterized fully in 3D (Fig. 5B; green circles) were classified as head-centered. In marked contrast, rFN cells were broadly distributed across categories with 52% of cells classified as either intermediate or body-centered. Consequently, when examined fully in 3D, NU cells could not account for the spatial tuning properties observed across the rFN population.

Importantly, although NU spatial tuning properties were generally inconsistent with transforming otolith signals towards body-centered coordinates, this does not necessarily rule out a role for these cells in the required transformations. In particular, theoretical studies have shown that cells at intermediate stages of neural networks involved in computing reference frame transformations often reflect posture-dependent changes in response gain (i.e., “gain fields”; Zipser and Andersen, 1988; Salinas and Abbott, 1995, 1996; Pouget and Sejnowski, 1997; Xing and Andersen, 2000; Deneve et al., 2001; Blohm et al., 2009). Thus, as a next step we also examined how cell response gains and phases in the cell’s PD changed across head orientations (Fig. 6A).

**Figure 6.**
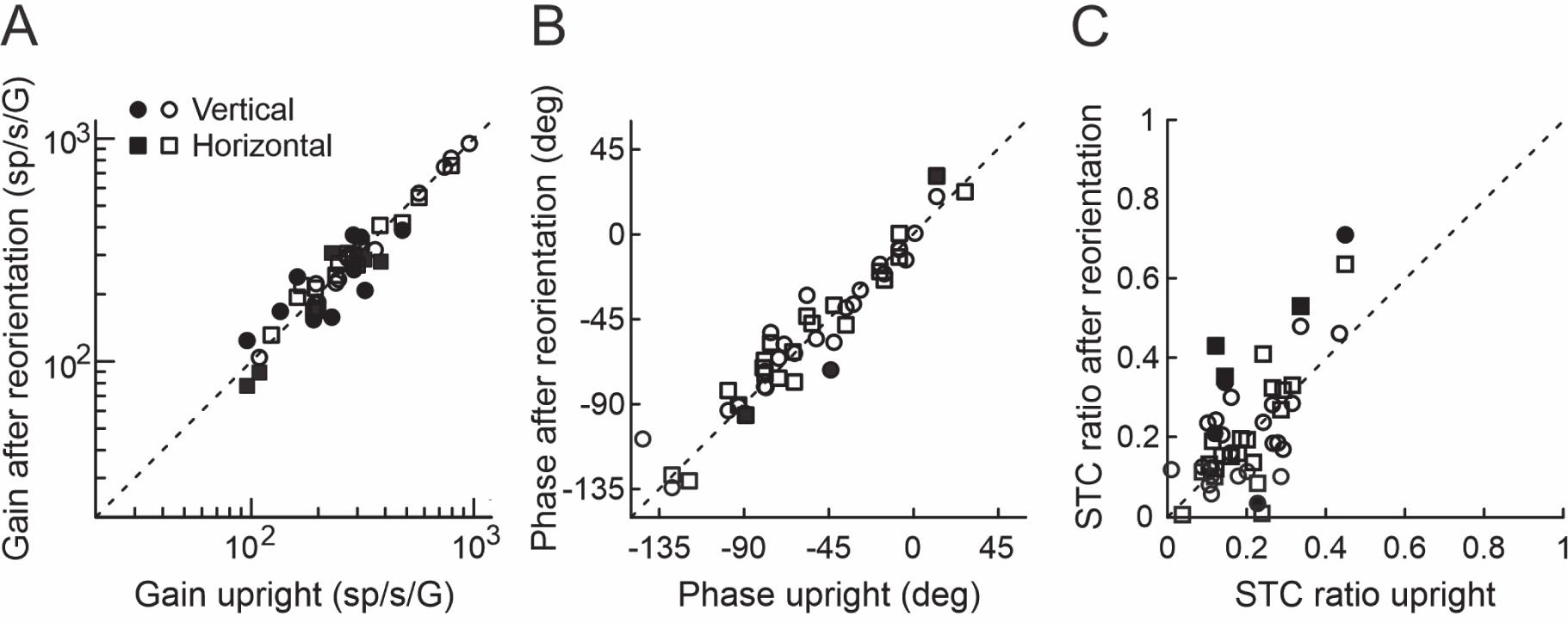
Dependence of translation tuning parameters on head orientation. Tuning parameters after head reorientation relative to the body in the vertical (circles) and horizontal (squares) planes are plotted versus the corresponding values with the head upright and straight forward. A, Gain in the maximum response direction. B, Phase in the maximum response direction. C, STC ratio. Filled symbols indicate significant differences after head reorientation as compared to upright/forward. Dashed line: unity slope.

We found that many cells exhibited changes in response gain that reached significance in 44% of cells (bootstrap analysis, 95% CIs on changes did not include 0). However, these changes were typically very small (average 13% change from upright or 0.32% change per degree of change in head angle). Furthermore, while comparable in size to the gain changes previously reported in the rFN and VN (0.36%/° in rFN and 0.24%/° in VN; Martin et al., 2018) they were substantially smaller than the posture-dependent gain-fields observed in cortical areas that have been implicated in reference frame transformations (e.g., ≈1%/° eye position in LIP and 1%/° - 3%/° in area 7a, Andersen et al., 1985; Andersen et al., 1990; 1.7%/° head/body-re-world position in area 7a, Snyder et al., 1998; 2%/° hand position and 3.4%/° eye position in PRR, Chang et al., 2009). No correlation was found between DI and the extent of gain change for reorientation in either horizontal or vertical planes (horizontal: r=0.12, p=0.58; vertical: r=-0.02, p=0.94). Other tuning properties varied little across head orientations, with a minority of cells showing significant changes in response phase (6.25%, Fig. 6B) or in the ratio of minimum to maximum response gain (STC ratio; 18.8%, Fig. 6C). Consequently, NU cells not only exhibited spatial tuning consistent with head-centered encoding of translation but showed little evidence for head-orientation-dependent changes in activity that might be expected at intermediate stages of a reference frame transformation.

### Spatial transformation of canal signals

The above results provide evidence that, like otolith afferents, NU cells encode otolith signals predominantly in a head-centered reference frame. However, unlike otolith afferents, which respond similarly to tilts and translations, most NU cells preferentially encode either translation or tilt (e.g., Fig 2A). Previous studies have shown that NU Purkinje cells reflect a solution to the tilt/translation ambiguity obtained by combining otolith signals with canal signals that have been spatially transformed into estimates of tilt relative to gravity (Yakusheva et al., 2007; Laurens et al., 2013). That is, in agreement with theoretical predictions (e.g., Green and Angelaki, 2004, 2007), canal signals in the NU are not head-centered but instead have been transformed into an estimate of the earth-horizontal component of rotation. However, whether such canal-derived tilt estimates are encoded in head- vs. body-centered coordinates remains unknown. Importantly, to reliably estimate translation, canal-derived tilt estimates must be appropriately matched to otolith signals across horizontal-plane changes in head orientation with respect to the body.

To confirm that this was indeed the case for NU translation-selective neurons, as well as to investigate the reference frames in which tilt-selective neurons encode motion, we characterized the spatial tuning of canal signals for a subset of recorded NU cells (30/95) across changes in body and/or head orientation. Cells were examined not only during translations but also during tilts and additive/subtractive combinations of tilt and translation stimuli (“Tilt-Translation” and “Tilt+Translation”; see Materials and Methods, Fig 1E). Among these, the Tilt-Translation stimulus is of particular relevance because it produces a net GIA of zero along the motion axis and thus no dynamic activation of otolith afferents tuned for that axis (Angelaki et al., 2004). This stimulus thus provides a means of isolating semicircular canal contributions to the cell’s response. To confirm that, as previously reported (Yakusheva et al., 2007; Laurens et al., 2013), the canal signals carried by NU cells had been spatially transformed into true 3D estimates of tilt (i.e., as opposed to reflecting head-centered rotation signals), cell responses were characterized for earth-vertical-axis rotations (EVAR) and for combinations of tilt and translation stimuli both with the head/body upright as well as after the whole-body was statically reoriented by 45° relative to gravity. Importantly, we compared cell responses to tilt and translation stimuli along the axis closest to the cell’s PD for Tilt-Translation with the body upright to those along the same axis after the body was statically reoriented along a perpendicular direction (e.g., body reorientation along the y-axis for a Tilt-Translation PD close to the x-axis; Fig 1Fi,ii). Consequently, the dynamic linear acceleration stimulus to the otoliths was similar across body orientations. In contrast, tilts stimulated different sets of canals in the different body orientations (i.e., mainly vertical canals with the body upright but a combination of vertical and horizontal canals after whole-body reorientation). Despite the different patterns of canal activation, cells carrying canal signals that had been transformed into true 3D tilt estimates should respond in the same way to tilt and combined tilt/translation stimuli across body orientations and remain unresponsive to earth-vertical-axis rotations (EVAR). Furthermore, cells encoding such canal-derived tilt estimates in head-centered coordinates should reflect spatial tuning for the Tilt-Translation stimulus (i.e., which isolates canal signals during earth-horizontal-axis rotation) that shifts with the head when it is reoriented relative to the body in the horizontal plane (Fig. 1Fiii).

Figure 7 shows an example translation-selective NU cell whose responses to translation, tilt and combined tilt/translation stimuli were characterized when upright as well as after whole-body reorientation relative to gravity and head-re-body reorientation in the horizontal plane. In the upright orientation, this cell responded robustly to translation but exhibited little response to tilt (Fig. 7Ai). When examined across horizontal-plane directions the cell’s translation response was largest for motions close to the 0° azimuth axis (Fig. 7C, left; blue trace). Consistent with this cell’s selectivity for translation, responses during Tilt+Translation and Tilt-Translation stimuli were similar to those during translation (Fig 7Ai). Furthermore, the spatial tuning and temporal properties of Tilt-Translation responses (Fig. 7C, right; blue trace) were closely aligned with those during translation, illustrating that the earth-horizontal canal and otolith components of the cell’s response were matched in the appropriate fashion to distinguish tilts and translations. No response was observed during rotations about an earth-vertical axis (Fig. 7Bi).

**Figure 7.**
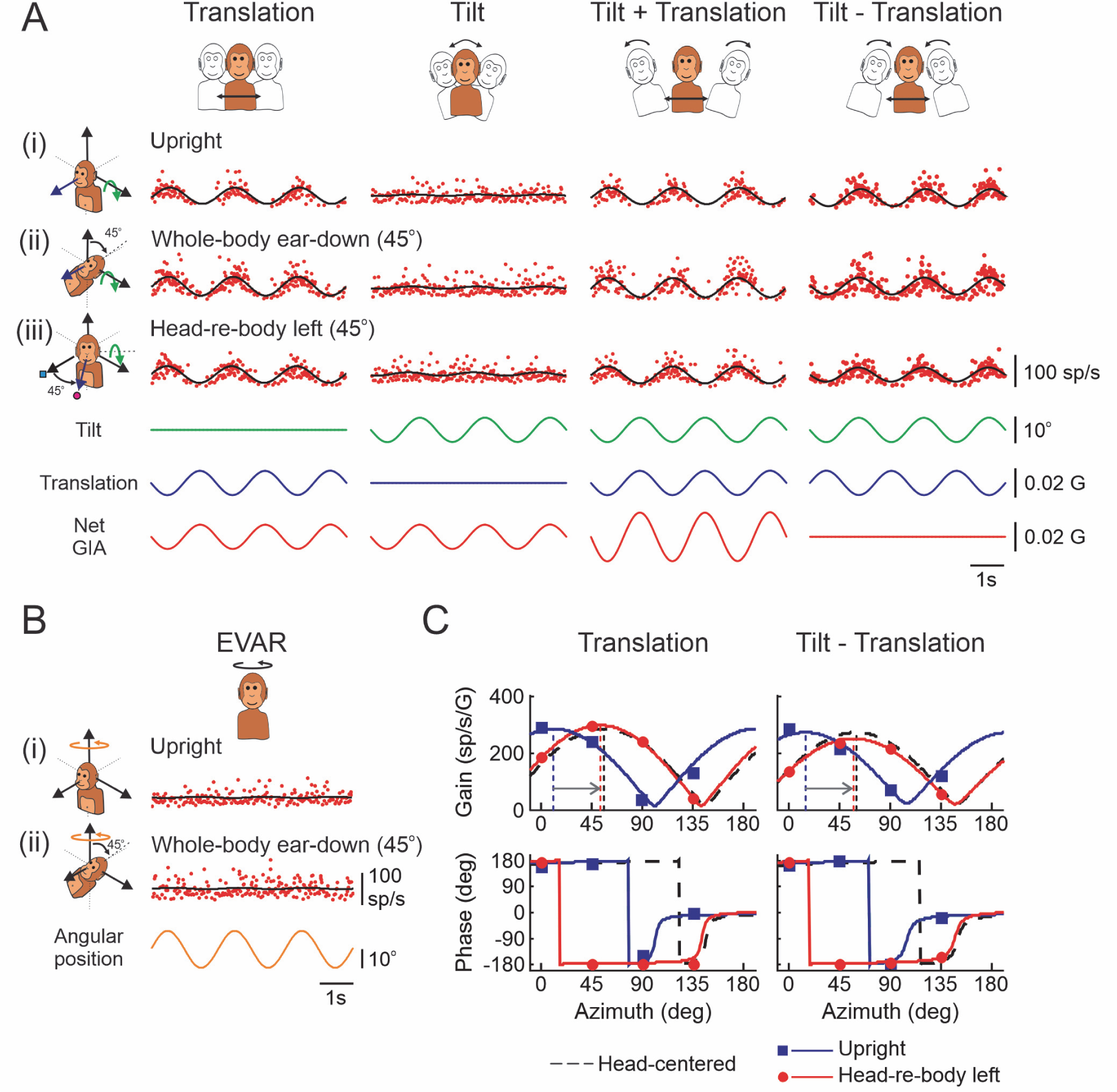
Example translation-encoding NU Purkinje cell reflecting a canal-derived estimate of tilt encoded in head-centered coordinates. A, IFR during Translation, Tilt, Tilt+Translation, and Tilt-Translation stimuli with the head upright (i), after whole-body reorientation by 45° toward ear-down (ii) and after head-re-body reorientation in horizontal plane by 45° to the left (iii). Illustrated responses are for tilt and translation stimuli applied along the head x-axis, which is aligned with the 0o azimuth direction in body coordinates in i,ii and the 45° azimuth direction in body coordinates after the head is reoriented relative to the body in iii. Black traces represent sinusoidal fits to IFRs. The stimulus traces (bottom) show tilt angular position (green), translational acceleration (blue) and net GIA (red). B, IFR responses during EVAR with the head upright (i) and after whole-body reorientation by 45o toward ear-down (ii). The stimulus traces (bottom) show earth-vertical angular position (orange). C, STC model fits to response gains and phases across horizontal-plane azimuth angles for Translation (left) and Tilt-Translation (right) stimuli with the head/body upright and forward (blue lines and symbols) and after head-re-body reorientation by 45° to the left (red lines and symbols). Gains and phases are expressed relative to translational acceleration. Dashed black curves in C indicate the predictions for head-centered tuning.

Similar observations were made after whole-body reorientation relative to gravity by 45° towards left-ear-down (i.e., along the 90° azimuth or y-axis). The cell remained selectively responsive to translation (Fig. 7Aii), exhibiting responses to tilts and translations along the 0° azimuth direction (i.e., x-axis) that were similar to those in the upright orientation. Furthermore, it responded robustly to Tilt-Translation motion but remained unresponsive to EVAR (Fig. 7Bii) even though this stimulus now activated the vertical canals – a canal stimulus to which the cell had responded robustly during upright Tilt-Translation motion (e.g., Fig. 7Ai, 4th column). These observations confirm that the cell’s response did not simply reflect head-centered canal signals but instead canal signals that had been spatially transformed into estimates of the earth-horizontal component of rotation (i.e., tilt), as required to resolve the tilt/translation ambiguity in 3D. Most importantly, when the head was reoriented relative to the body by 45° in the horizontal plane (Fig. 7Aiii), not only did the cell’s PDs for translation shift with the head by 42° (DI of 0.93) towards the prediction for head-centered coding (Fig. 7C, left, red trace closely aligned with black dashed trace) but the cell’s PD for Tilt-Translation shifted to a similar extent (43°; DI of 0.96; Fig. 7C, right). Neither DI was statistically different from 1 (bootstrap test; 95% CIs included 1). Consequently, this cell encoded both otolith signals and canal-derived tilt signals in a head-centered reference frame.

Similar conclusions were reached for tilt-selective neurons as illustrated for the example cell in Figure 8. In contrast to the translation-encoding neuron of Figure 7, this cell responded robustly to all stimuli involving tilt but exhibited little response to pure translation (Fig. 8Ai). Like the cell of Figure 7, however, this tilt-selective cell exhibited little response to EVAR both upright and after whole-body reorientation (Fig. 8B). Furthermore, its responses to tilt remained consistent after whole-body reorientation (Fig. 8Aii), revealing that it did not simply encode head-centered vertical canal signals but instead a combination of vertical and horizontal canal-derived signals that had been spatially transformed into an estimate of reorientation relative to gravity. Most importantly, when the head was reoriented in the horizontal plane (Fig. 8Aiii) the cell’s spatial tuning for tilt and Tilt-Translation stimuli shifted with the head (Fig. 8C) consistent with the predictions for head-centered tilt coding (Tilt DI: 0.99; Tilt-Translation DI: 0.94).

**Figure 8.**
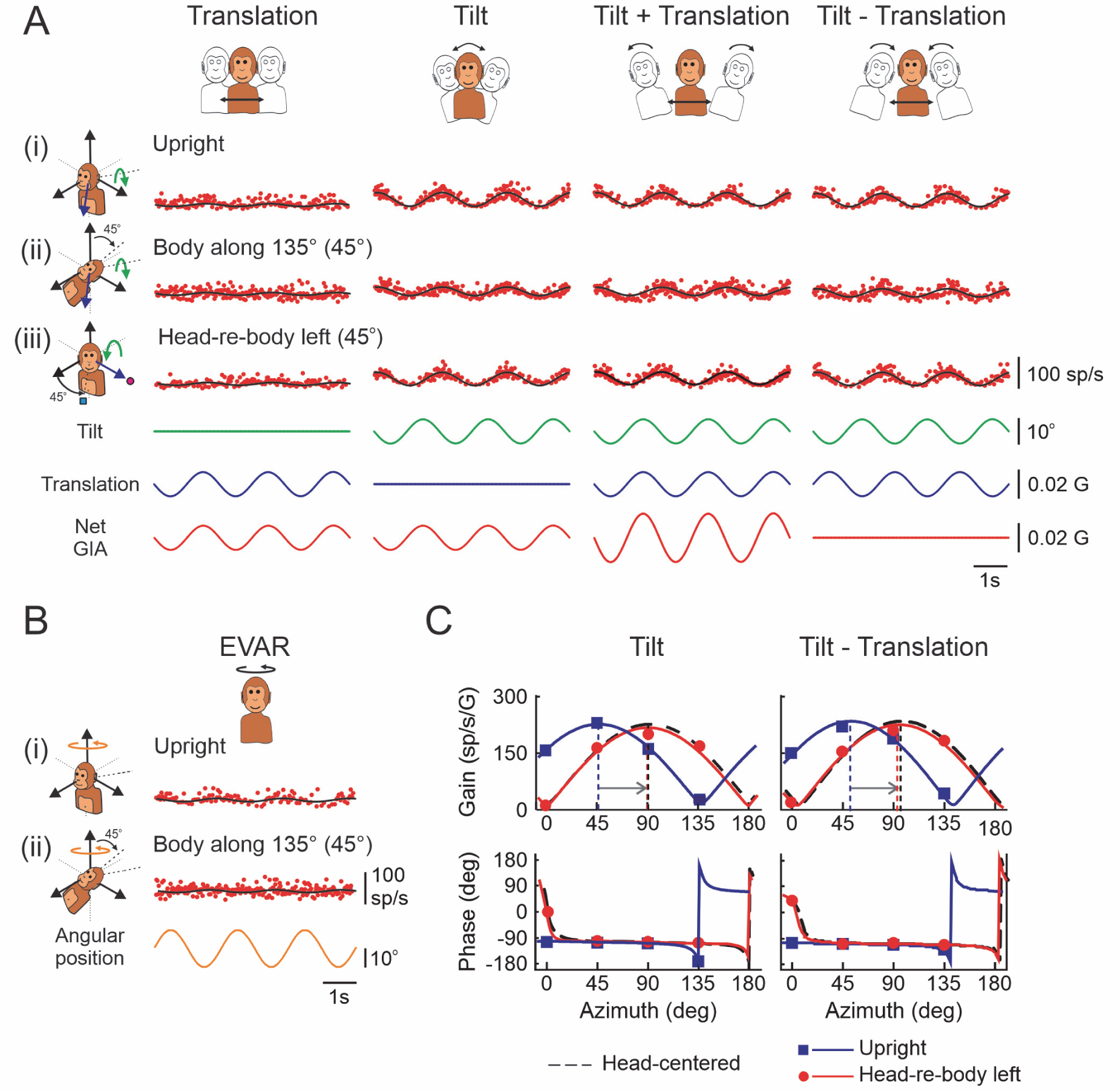
Example tilt-encoding NU Purkinje cell. A, IFR responses during Translation, Tilt, Tilt+Translation, and Tilt-Translation stimuli with the head upright (i), after whole-body reorientation by 45° along the 135° azimuth direction (ii) and after head-re-body reorientation in horizontal plane by 45° to the left (iii). Illustrated responses are for tilt and translation stimuli applied along the head 45° azimuth axis which is aligned with the body 45o azimuth direction in i,ii and the 90° body azimuth direction after the head is reoriented relative to the body in iii. Black traces represent sinusoidal fits to IFRs. The stimulus traces (bottom) show tilt angular position (green), translational acceleration (blue) and net GIA (red). B, IFR responses during EVAR with the head upright (i) and after whole-body reorientation by 45° along the 135° azimuth direction (ii). The stimulus traces (bottom) show angular position (orange). C, STC model fits to response gains and phases across horizontal-plane azimuth angles for Tilt (left) and Tilt-Translation (right) stimuli with the head/body upright and forward (blue lines and symbols) and after head-re-body reorientation by 45° to the left (red lines and symbols). Gains and phases are expressed relative to the acceleration due to tilt. Dashed black curves in C indicate the predictions for head-centered tuning.

Figure 9 summarizes the responses observed for the subset of NU cells examined for different combinations of tilt and translation motions across changes in whole-body orientation (N=22). As shown in Figure 9A, response gains for the different motion stimuli were similar across whole-body orientations, reflecting population regression slopes across stimuli that were close to and not statistically different from one for otolith and canal response components considered individually (translation slope: 0.96, CI: [0.88,1.05]; Tilt-Translation slope: 0.89, CI: [0.72,1.05]; all stimuli slope: 0.91, CI: [0.86,0.97]). Furthermore, for translation-sensitive neurons otolith and canal-derived tilt signals remained appropriately amplitude-matched (i.e., as required to estimate translation) as reflected in ratios of their response gains during Tilt-Translation as compared to translation (Tilt-Trans/Trans) that were distributed close to 1 both upright and after whole-body reorientation (mean upright: 0.89; body reoriented: 0.94).

**Figure 9.**
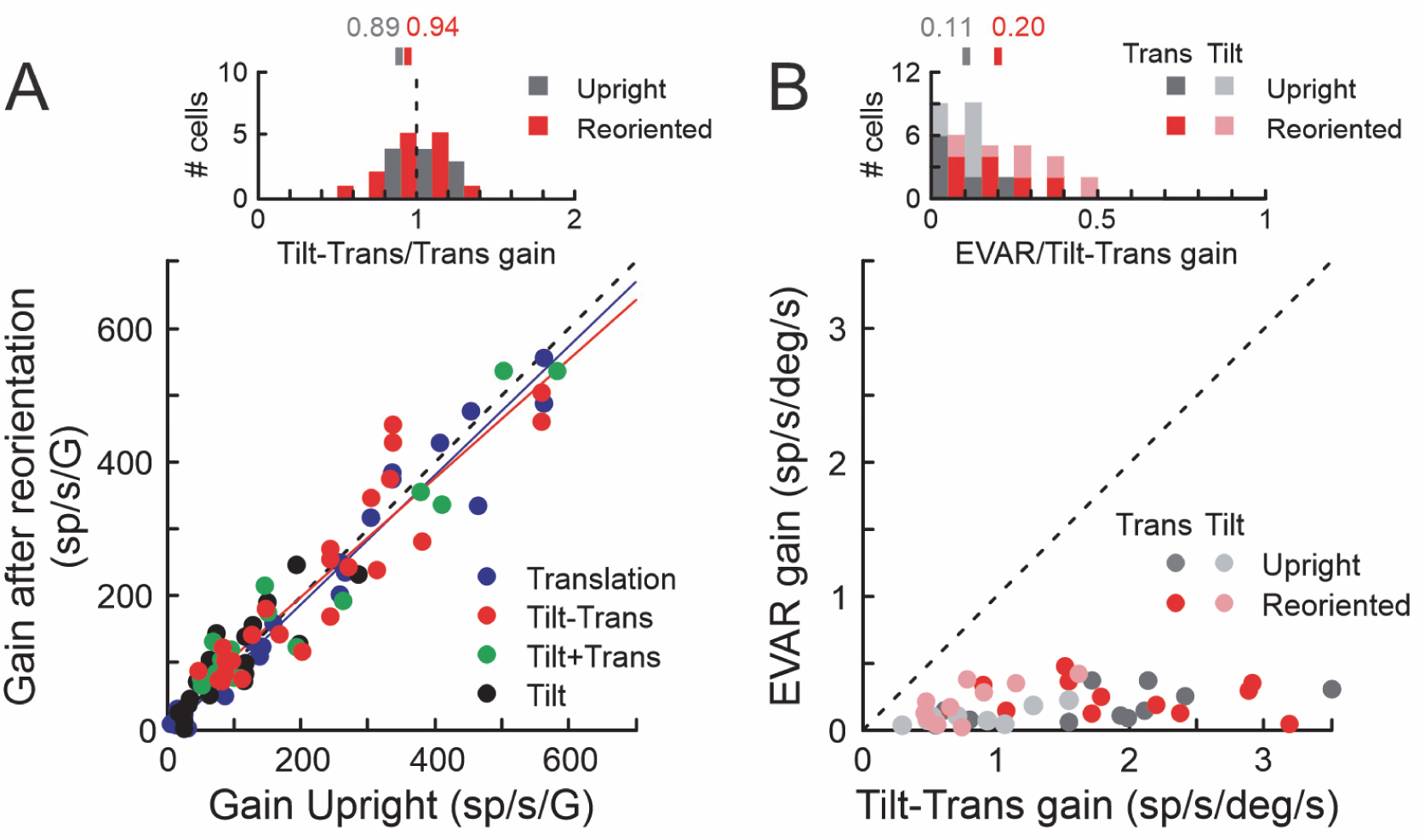
Evidence that NU cells carry canal-derived tilt estimates. **A**, Response gains after whole-body reorientation relative to the gravity for Translation (blue), Tilt (black), Tilt+Translation (green) and Tilt-Translation (red) stimuli plotted versus the corresponding gains with the whole-body upright (N=25 reorientations across 22 cells). Response gains in both body orientations are for motions along the axis (0°, 45°, 90°, 135°) closest to the cell’s upright horizontal-plane PD for the Tilt-Translation stimulus. The whole-body was reoriented by 45° along the axis perpendicular to this PD direction (e.g., see Fig 1Fi,ii). The top inset shows the distribution of Tilt-Translation/Translation gain ratios with the head upright (gray) as compared to after body reorientation (red) for all translation-sensitive cells (i.e., all except tilt cells; N=11 cells and 14 reorientations). Vertical bars above the plot indicate distribution means. **B**, Cell response gain for canal stimulation during earth-vertical-axis rotation (EVAR) plotted versus canal response gain for earth-horizontal-axis rotation during Tilt-Translation motion with the head upright (gray; N=20) and after whole-body reorientation (red; N=22 reorientations across 20 cells). The Tilt-Translation motion direction and body reorientation axis for each cell were the same as in **A**. Top inset shows the distribution of EVAR/Tilt-Translation canal gain ratios with the head upright (gray) as compared to after body reorientation (red). Vertical bars above the plot indicate distribution means. Dark gray/red: translation cells; Pale gray/red: tilt cells.

More direct evidence that recorded NU cells carried a canal-derived tilt estimate was obtained by comparing their responses to canal stimulation during earth-vertical-axis rotations to those during earth-horizontal-axis rotations using the Tilt-Translation stimulus (Fig. 9B). Cells reflecting such a tilt estimate should respond during Tilt-Translation motion but should not respond to EVAR. Consistent with this, in the upright orientation NU cells exhibited negligible responses to EVAR which stimulated mainly the horizontal canals, but substantial responses to Tilt-Translation motion which stimulated mainly the vertical canals (Fig. 9B; gray circles). This was reflected in a mean EVAR/Tilt-Trans ratio of 0.11 (Fig. 9B inset, gray). Similarly, after whole-body reorientation cells continued to reflect negligible responses to EVAR even though in the new body orientation EVAR now activated both vertical and horizontal canals (Fig. 9B; red circles). Importantly, after whole-body reorientation by 45° both EVAR and Tilt-Translation motion activated the vertical (and horizontal) canals to similar extents. Thus, if the responses to Tilt-Translation motion observed when upright were due to head-centered vertical canal signals, after whole-body reorientation responses to EVAR and Tilt-Translation motion should have been similar in amplitude (i.e., EVAR/Tilt-Trans ratio close to 1). In contrast, EVAR/Tilt-Trans ratios after reorientation remained close to zero (mean EVAR/Tilt-Trans of 0.20) consistent with the transformation of canal signals on these cells into true estimates of dynamic tilt.

Finally, to investigate whether NU cells encoded canal-derived tilt signals in head- versus body-centered coordinates, their spatial tuning in the body upright orientation with the head facing forward was compared to that after head reorientation to the left. As illustrated in Figure 10A, DIs for responses to Tilt-Translation motion (i.e., canal-derived estimates of tilt) were narrowly distributed about 1 consistent with head-centered coding (median DI of 1.01 not different from 1; Wilcoxon signed rank test, p=0.96). Furthermore, model fits across horizontal plane head orientations revealed that cell responses to Tilt-Translation motion were significantly better correlated with a head-centered as compared to a body-centered model (Fig. 10B). Consequently, like otolith signals, canal-derived estimates of tilt relative to gravity were encoded in head-centered coordinates.

**Figure 10.**
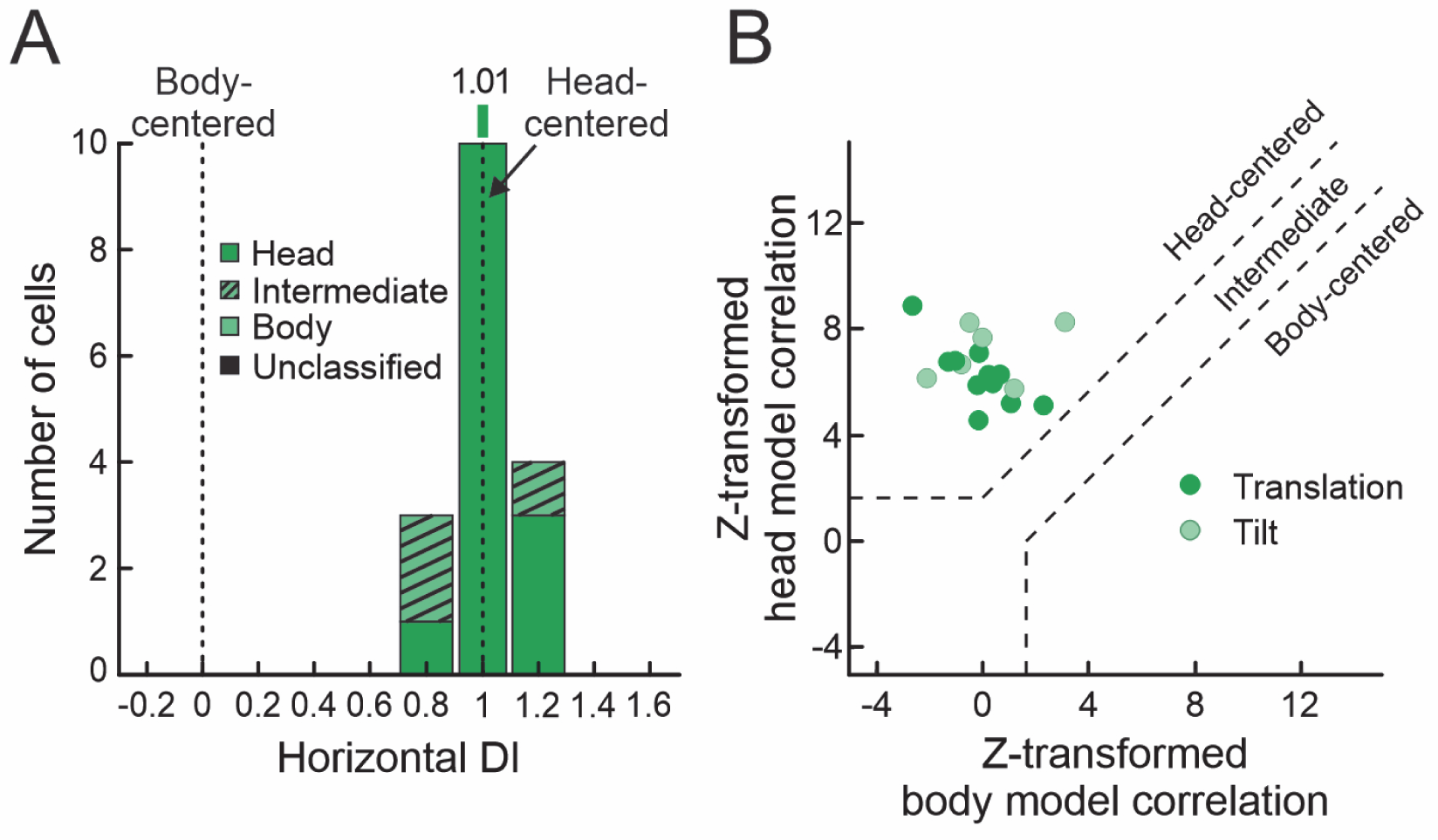
Head-centered coding of canal-derived tilt signals. **A**, Distribution of DI values for the canal-derived tilt component of NU cell responses (isolated during Translation-Tilt motion) across changes in head orientation in the horizontal plane (N=17, including 11 translation and 6 tilt cells). DIs were classified based on bootstrapped DI 95% CIs as head-centered (dark colors), body-centered (pale colors) or intermediate (hatched). The vertical bar above the plot indicates the population median. **B**, Correlation of NU canal-derived tilt signals (i.e., during Tilt-Translation motion) with head- and body-centered models. The plot shows partial correlation coefficients (Z-transformed) for head-centered versus body-centered model fits across two head-re-body orientations (upright and after horizontal plane reorientation). Significance regions are based on the difference between head- and body-centered model fit Z-scores corresponding to p<0.05 (top left: head-centered; bottom right: body-centered; middle diagonal region: not significantly better correlated with either model and classified as “intermediate”). Dark green: translation cells; Pale green: tilt cells.

## Discussion

Daily activities such as postural control (Maurer et al., 2006; Karmali et al., 2021; Mildren and Cullen, 2023), navigation (Yoder and Taube, 2014), spatial perception (Clemens et al., 2011; Gu, 2018) and voluntary movement (Moreau-Debord et al., 2014; Blouin et al., 2015; Keyser et al., 2017; Martin et al., 2021) all rely on estimates of our body’s motion and orientation in space. Vestibular signals are essential for such estimates, but to contribute appropriately, the motions associated with each sensory signal must be correctly interpreted and they must be transformed into body-centered coordinates.

Here, we characterized for the first time the extent to which the nodulus and uvula of the posterior vermis – regions implicated in resolving the otolith tilt/translation ambiguity - show evidence for body-centered coding of motion. In contrast to neurons downstream in the rFN, many of which reflect a transformation of vestibular signals towards body-centered coordinates (Kleine et al., 2004; Shaikh et al., 2004; Martin et al., 2018), we found that the 3D tuning properties of NU cells were consistent with head-centered coding of otolith signals. Furthermore, in keeping with the computations necessary to resolve the tilt/translation ambiguity (Merfeld et al., 1993; Bos and Bles, 2002; Zupan et al., 2002; Green and Angelaki, 2004, 2007; Laurens and Angelaki, 2017) canals signals in the NU were not head-centered, but instead had been spatially transformed into world- (or gravity-) referenced estimates of tilt (Yakusheva et al., 2007; Laurens et al., 2013). Notably, however, the resulting tilt estimates shifted with changes in head-re-body position in the horizontal plane indicating that they are encoded in head-centered coordinates. These results thus imply that the body-centered estimates of translation and tilt required for many of our daily behaviors are computed elsewhere, either by further transformation of NU outputs or via parallel computations in other regions.

### Estimates of translation and tilt in the NU

Compatible with previous investigations (Yakusheva et al., 2007; Laurens et al., 2013; Laurens and Angelaki, 2020) Purkinje cells in the primate NU could be categorized into functional groups that selectively encoded either translation or tilt, with only a minority (10%) having responses that were not significantly better correlated with either of the two motions or reflected the net GIA. Furthermore, as shown previously (Yakusheva et al., 2007) cells in this region exhibited little response to earth-vertical-axis rotations, regardless of body-re-gravity orientation, but responded robustly to canal stimulation during earth-horizontal-axis rotation from upright (i.e., Tilt-Translation motion). Here, we further show that such canal-derived responses to earth-horizontal-axis rotation remain consistent across substantial changes in body-re-gravity orientation, thus explicitly confirming that both vertical and horizontal canal signals have been combined and spatially transformed on these cells into estimates of tilt with respect to gravity.

Previous work by Laurens et al. (2013) was the first to reveal true tilt-encoding cells in the NU by showing that these cells responded similarly to sinusoidal tilts from upright and to tilt produced by constant velocity yaw rotation about an axis tilted 10° from upright (i.e., off-vertical-axis rotation, OVAR). Importantly, the stimuli used by Laurens et al. (2013), combined a constant velocity stimulus to the horizontal canals with a cyclic stimulus to the otoliths (i.e., GIA signal) to provide evidence for the 3D computations required to estimate tilt relative to gravity. The stimuli used in the current experiments complement and extend these findings by explicitly isolating the canal signal contributions to these computations (i.e., using the Tilt-Translation stimulus) and confirming not only that they have been spatially transformed into estimates of tilt, but that canal and otolith signals remain appropriately matched to resolve the tilt/translation ambiguity over a broad range of body orientations (i.e., up to at least 45° reorientation from upright). Collectively, our observations thus support previous conclusions that NU Purkinje cells reflect the output of the 3D computations necessary for tilt/translation discrimination (Yakusheva et al., 2007; Laurens et al., 2013).

Most importantly, however, we extend previous results to show that translation and tilt estimates in the NU are encoded in head-centered coordinates. This finding also emphasizes a critical distinction in the reference frames associated with what representations NU cells encode (i.e., translation and tilt) versus those of the sensory signals (i.e., otolith and canal) used to compute them. Specifically, to resolve the tilt/translation ambiguity canal signals in the NU must be spatially transformed into an estimate of the earth-horizontal component of rotation that signals dynamic tilt. Consequently, canal signals in the NU are not head-centered but rather reflect world/gravity-referenced coding of rotation. However, the dynamic tilt estimates that result from those transformations are head-centered - they exhibit tuning that remains fixed to a particular head axis across changes in static head-re-body orientation. This further emphasizes that NU tilt cells specifically encode signals relevant for estimating head (but not body) orientation with respect to gravity.

### Distinct pathways for computing head- vs body-centered motion estimates?

If motion estimates in the NU are encoded predominantly in head-centered coordinates, then where are body-centered tilt and translation estimates computed? Because many rFN neurons reflect a transformation towards body-centered coordinates (Kleine et al., 2004; Shaikh et al., 2004; Martin et al., 2018) one obvious answer is that the transformation occurs downstream of the NU in the rFN. Yet, there is evidence to suggest that at least a portion of the computation may occur elsewhere, in regions of anterior vermis (AV) that project to the rFN. Previous findings implicate these regions in transforming vestibular signals towards body-centered coordinates (Manzoni et al., 1998; Manzoni et al., 1999). Furthermore, dynamic neck proprioceptive and vestibular signals combine on many AV Purkinje cells to encode representations intermediate between head and body motion (Zobeiri and Cullen, 2022). Both observations support a role for the AV in computing body-centered motion representations. This is also compatible with our recent suggestion that the bulk of the non-linear computations to effect the head-to-body transformation occur upstream of the rFN in the cerebellar cortex (Martin et al., 2018). Here we have shown that they do not occur in the NU, suggesting that they may instead occur either in the AV or in a distributed AV-rFN circuit. But if the computations to effect the head-to-body reference frame transformation take place largely upstream in the AV how then do rFN cells also inherit an at least partial solution to the tilt/translation ambiguity?

Although it has been suggested that a solution to the tilt/translation ambiguity may be conveyed to the rFN via direct projections from the NU (e.g., Angelaki et al., 2010) there are other possibilities. One is that tilt and translation signals from the NU are conveyed only indirectly to the rFN via brainstem circuits and the AV (Ruggiero et al., 1977; Kotchabhakdi and Walberg, 1978; Barmack, 2003; Lee et al., 2014) where such estimates are further transformed and integrated with proprioceptive signals. Another intriguing possibility is that a solution to the tilt/translation ambiguity is independently computed in the AV itself. Both possibilities are compatible with anatomical studies showing that the most rostral portions of the FN, strongly implicated in body postural control and locomotion, receive abundant projections from the AV (Haines, 1976; Armstrong and Schild, 1978; Fujita et al., 2020). In contrast, the NU projects primarily to ventral regions of the FN including relatively caudal as well as slightly more rostral regions (Haines, 1977; Armstrong and Schild, 1978; Dietrichs, 1983; Fujita et al., 2020) that, at least in rodents, appear to target populations largely distinct from those receiving abundant projections from the AV (Fujita et al., 2020). Because the neurons recorded in the study of Martin et al., (2018) (e.g., Fig. 4) were broadly distributed across the rFN, it seems likely that a substantial subset of them received projections from the AV. However, it remains unclear to what extent motion-sensitive neurons either in that study or in other studies of the primate rFN (e.g., Siebold et al., 1997; Zhou et al., 2001; Kleine et al., 2004; Shaikh et al., 2004; Mackrous et al., 2019) corresponded to populations receiving NU projections.

Vestibular processing through at least partially distinct cerebellar cortical pathways may also explain other differences in the motion representations found in the rFN versus NU. In particular, whereas a majority of NU cells can be classified as predominantly tilt or translation-coding, rFN cell properties are more broadly distributed with many cells responding substantially to both translation and tilt (Angelaki et al., 2004; Green et al., 2005; Laurens and Angelaki, 2016; Mackrous et al., 2019) and/or to earth-vertical-axis rotations (Siebold et al., 1997; Siebold et al., 2001; Zhou et al., 2001; Shaikh et al., 2005a). These more complex properties may reflect encoding of specific types of body motion representations for different behaviors (e.g., trunk postural control, reaching, locomotion) where estimates of pure translation or tilt about head-centered axes may be less relevant. Such representations could arise in the rFN simply by combining tilt and translation computed in the NU with other more afferent-like vestibular signals (e.g., conveyed via mossy fibre collaterals; Ruggiero et al., 1977). Alternatively, however, they might reflect vestibular processing through different cerebellar cortical pathways (e.g., through the AV). Future work that examines the evidence for a solution to the tilt/translation ambiguity in the AV as well as the reference frames (i.e., head vs body vs world-centered) in which both otolith and canal signals are encoded will aid in distinguishing these possibilities. Likewise, although rFN neurons have been identified that preferentially encode either translation or tilt from upright, the extent to which cells with different characteristics truly encode allocentric tilt versus egocentric rotation signals remains to be established (e.g., by testing across different body orientations; but see Buron et al., 2022).

Despite many unanswered questions, our current findings, together with the observations summarized above, lead us to suggest that there may exist largely distinct cerebellar pathways or modules for computing head- versus body-centered motion representations, each of which may reflect at least a partial solution to the tilt/translation ambiguity and subserve distinct functional roles. In particular, our finding that translation and tilt estimates in the NU are predominantly head-centered suggests a primary role for these signals in stabilizing the head and gaze in space. This is consistent with direct NU projections to portions of the vestibular nuclei from which the medial vestibulo-spinal tract (i.e., mediating vestibulocollic reflexes) originates as well as to brainstem regions involved in the control of eye movements (Walberg and Dietrichs, 1988; Barmack, 2003; Meng et al., 2014). Pathways through the NU may thus be integral to forming a stable earth-horizontal platform for integrating visual and vestibular cues that serves as a reference for perception of our motion and orientation relative to the external world, essential for both motor (e.g., postural control; Massion, 1994; Mergner and Rosemeier, 1998) and more cognitive functions (e.g., spatial orientation for navigation; Rochefort et al., 2013; Yoder and Taube, 2014; Laurens et al., 2016). Consistent with this proposal, NU lesions not only give rise to substantial body and head postural instability (Dow, 1938; Moon et al., 2009) but more specifically disrupt the influence of gravity on spatial orientation of vestibular-induced eye movements (Angelaki and Hess, 1995; Wearne et al., 1998) and perception of visual earth-vertical (Tarnutzer et al., 2015).

In contrast, pathways through the AV and rFN, contributing to body-centered motion estimates, may play a more direct role in online corrections mediated by axial and limb muscles to maintain postural equilibrium during stance and gait (Massion, 1994; Horak and Macpherson, 1996; Peterka, 2002), as well as in estimating body motion for purposes such as heading perception (Gu, 2018), path integration (Rondi-Reig et al., 2022) and goal-directed action (Medendorp and Selen, 2017; Martin et al., 2021). This is consistent with abundant rFN projections to brainstem regions involved in body postural control and locomotion (Batton et al., 1977; Homma et al., 1995; Fujita et al., 2020), as well as to regions of the thalamus that project to parietal areas implicated in encoding body-centered motion information (Akbarian et al., 1992; Meng et al., 2007; Chen et al., 2013). Future work that further explores the nature of the motion representations within different cerebellar pathways and their downstream targets will be important to elucidate their functional contributions to a broad range of behaviors.

## Acknowledgements

This work was supported by an operating grant from the Canadian Institutes of Health Research (CIHR: PJT-153257), an infrastructure grant from the Canadian Foundation for Innovation (CFI) as well as a CIHR PhD fellowship to CM and CIHR MSc fellowship to FB. We thank P. Cisek for comments on the manuscript and M. Latourelle for excellent technical assistance.

## References

Akbarian S, Grusser OJ, Guldin WO (1992) Thalamic connections of the vestibular cortical fields in the squirrel monkey (Saimiri sciureus). The Journal of comparative neurology 326:423–441.

Andersen RA, Essick GK, Siegel RM (1985) Encoding of spatial location by posterior parietal neurons. Science, NY 230:456–458.

Andersen RA, Bracewell RM, Barash S, Gnadt JW, Fogassi L (1990) Eye position effects on visual, memory, and saccade-related activity in areas LIP and 7a of macaque. J Neurosci 10:1176–1196.

Angelaki DE (1991) Dynamic polarization vector of spatially tuned neurons. IEEE transactions on bio-medical engineering 38:1053–1060.

Angelaki DE, Hess BJ (1995) Inertial representation of angular motion in the vestibular system of rhesus monkeys. II. Otolith-controlled transformation that depends on an intact cerebellar nodulus. Journal of neurophysiology 73:1729–1751.

Angelaki DE, Dickman JD (2000) Spatiotemporal processing of linear acceleration: primary afferent and central vestibular neuron responses. Journal of neurophysiology 84:2113–2132.

Angelaki DE, Shaikh AG, Green AM, Dickman JD (2004) Neurons compute internal models of the physical laws of motion. Nature 430:560–564.

Angelaki DE, Yakusheva TA, Green AM, Dickman JD, Blazquez PM (2010) Computation of egomotion in the macaque cerebellar vermis. Cerebellum (London, England) 9:174–182.

Armstrong DM, Schild RF (1978) An investigation of the cerebellar cortico-nuclear projections in the rat using an autoradiographic tracing method. I. Projections from the vermis. Brain research 141:1–19.

Barmack NH (2003) Central vestibular system: vestibular nuclei and posterior cerebellum. Brain research bulletin 60:511–541.

Barmack NH, Shojaku H (1995) Vestibular and visual climbing fiber signals evoked in the uvula-nodulus of the rabbit cerebellum by natural stimulation. Journal of neurophysiology 74:2573–2589.

Batton RR, 3rd, Jayaraman A, Ruggiero D, Carpenter MB (1977) Fastigial efferent projections in the monkey: an autoradiographic study. The Journal of comparative neurology 174:281–305.

Berens P (2009) CircStat: A matlab toolbox for circular statistics. Statistical Software 31.

Blohm G, Keith GP, Crawford JD (2009) Decoding the cortical transformations for visually guided reaching in 3D space. Cereb Cortex 19:1372–1393.

Blouin J, Bresciani JP, Guillaud E, Simoneau M (2015) Prediction in the Vestibular Control of Arm Movements. Multisensory research 28:487–505.

Bos JE, Bles W (2002) Theoretical considerations on canal-otolith interaction and an observer model. Biological cybernetics 86:191–207.

Brooks JX, Cullen KE (2009) Multimodal integration in rostral fastigial nucleus provides an estimate of body movement. J Neurosci 29:10499–10511.

Brooks JX, Cullen KE (2013) The primate cerebellum selectively encodes unexpected self-motion. Curr Biol 23:947–955.

Buron F, Martin CZ, Green AM (2022) Reference frames for encoding vestibular signals in the posterior vermis and rostral fastigial nucleus of the cerebellum across changes in head and body orientation. Society for Neuroscience Conference, San Diego, Program No. 632.08, Neuroscience Meeting Planner,Online.

Chang SW, Papadimitriou C, Snyder LH (2009) Using a compound gain field to compute a reach plan. Neuron 64:744–755.

Chen-Huang C, Peterson BW (2006) Three dimensional spatial-temporal convergence of otolith related signals in vestibular only neurons in squirrel monkeys. Experimental brain research 168:410–426.

Chen X, Deangelis GC, Angelaki DE (2013) Diverse spatial reference frames of vestibular signals in parietal cortex. Neuron 80:1310–1321.

Clemens IA, De Vrijer M, Selen LP, Van Gisbergen JA, Medendorp WP (2011) Multisensory processing in spatial orientation: an inverse probabilistic approach. J Neurosci 31:5365–5377.

Deneve S, Latham PE, Pouget A (2001) Efficient computation and cue integration with noisy population codes. Nature neuroscience 4:826–831.

Dietrichs E (1983) The cerebellar corticonuclear and nucleocortical projections in the cat as studied with anterograde and retrograde transport of horseradish peroxidase. V. The posterior lobe vermis and the flocculo-nodular lobe. Anatomy and embryology 167:449–462.

Dow RS (1938) Effect of lesions in the vestibular part of the cerebellum in primates Archives of Neurology & Psychiatry 40:500–520.

Efron B, Tibshirani RJ (1993) An Introduction to the Bootstrap. Boca Raton, FL: Chapman and Hall.

Fernandez C, Goldberg JM (1976a) Physiology of peripheral neurons innervating otolith organs of the squirrel monkey. III. Response dynamics. Journal of neurophysiology 39:996–1008.

Fernandez C, Goldberg JM (1976b) Physiology of peripheral neurons innervating otolith organs of the squirrel monkey. I. Response to static tilts and to long-duration centrifugal force. Journal of neurophysiology 39:970–984.

Fujita H, Kodama T, Du Lac S (2020) Modular output circuits of the fastigial nucleus for diverse motor and nonmotor functions of the cerebellar vermis. Elife 9:e58613.

Green AM, Angelaki DE (2004) An integrative neural network for detecting inertial motion and head orientation. Journal of neurophysiology 92:905–925.

Green AM, Angelaki DE (2007) Coordinate transformations and sensory integration in the detection of spatial orientation and self-motion: from models to experiments. Progress in brain research 165:155–180.

Green AM, Angelaki DE (2010) Multisensory integration: resolving sensory ambiguities to build novel representations. Current opinion in neurobiology 20:353–360.

Green AM, Shaikh AG, Angelaki DE (2005) Sensory vestibular contributions to constructing internal models of self-motion. Journal of neural engineering 2:S164–179.

Gu Y (2018) Vestibular signals in primate cortex for self-motion perception. Current opinion in neurobiology 52:10–17.

Haines DE (1976) Cerebellar corticonuclear and corticovestibular fibers of the anterior lobe vermis in a prosimian primate (Galago senegalensis). The Journal of comparative neurology 170:67–95.

Haines DE (1977) Cerebellar corticonuclear and corticovestibular fibers of the flocculonodular lobe in a prosimian primate (Galago senegalensis). The Journal of comparative neurology 174:607–630.

Hartigan JA, Hartigan PM (1985) The dip test of unimodality. Annals of Statistics 13:70–84.

Hernández RG, De Zeeuw CI, Zhang R, Yakusheva TA, Blazquez PM (2020) Translation information processing is regulated by protein kinase C-dependent mechanism in Purkinje cells in murine posterior vermis. Proceedings of the National Academy of Sciences 117:17348–17358.

Homma Y, Nonaka S, Matsuyama K, Mori S (1995) Fastigiofugal projection to the brainstem nuclei in the cat: an anterograde PHA-L tracing study. Neuroscience research 23:89–102.

Horak FB, Macpherson JM (1996) Postural orientation and equilibrium. In: Handbook of physiology, exercise: regulation and integration of multiple systems, pp 255–292. New York: Oxford University Press.

Karmali F, Goodworth AD, Valko Y, Leeder T, Peterka RJ, Merfeld DM (2021) The role of vestibular cues in postural sway. Journal of neurophysiology 125:672–686.

Keyser J, Medendorp WP, Selen LPJ (2017) Task-dependent vestibular feedback responses in reaching. Journal of neurophysiology 118:84–92.

Kleine JF, Guan Y, Kipiani E, Glonti L, Hoshi M, Buttner U (2004) Trunk position influences vestibular responses of fastigial nucleus neurons in the alert monkey. Journal of neurophysiology 91:2090–2100.

Kotchabhakdi N, Walberg F (1978) Cerebellar afferent projections from the vestibular nuclei in the cat: an experimental study with the method of retrograde axonal transport of horseradish peroxidase. Experimental brain research 31:591–604.

Laurens J, Angelaki DE (2016) How the vestibulocerebellum builds an internal model of self-motion. In: The Neuronal Codes of the Cerebellum, pp 97–115: Elsevier.

Laurens J, Angelaki DE (2017) A unified internal model theory to resolve the paradox of active versus passive self-motion sensation. Elife 6:e28074.

Laurens J, Angelaki DE (2020) Simple spike dynamics of Purkinje cells in the macaque vestibulo-cerebellum during passive whole-body self-motion. Proceedings of the National Academy of Sciences of the United States of America 117:3232–3238.

Laurens J, Meng H, Angelaki DE (2013) Neural representation of orientation relative to gravity in the macaque cerebellum. Neuron 80:1508–1518.

Laurens J, Kim B, Dickman JD, Angelaki DE (2016) Gravity orientation tuning in macaque anterior thalamus. Nature neuroscience 19:1566–1568.

Lee RX, Huang JJ, Huang C, Tsai ML, Yen CT (2014) Collateral projections from vestibular nuclear and inferior olivary neurons to lobules I/II and IX/X of the rat cerebellar vermis: a double retrograde labeling study. The European journal of neuroscience 40:2811–2821.

Liu S, Dickman JD, Angelaki DE (2011) Response dynamics and tilt versus translation discrimination in parietoinsular vestibular cortex. Cerebral Cortex 21:563–573.

Luan H, Gdowski MJ, Newlands SD, Gdowski GT (2013) Convergence of vestibular and neck proprioceptive sensory signals in the cerebellar interpositus. J Neurosci 33:1198–1210a.

Mackrous I, Carriot J, Jamali M, Cullen KE (2019) Cerebellar Prediction of the Dynamic Sensory Consequences of Gravity. Curr Biol 29:2698–2710 e2694.

Manzoni D, Pompeiano O, Andre P (1998) Neck influences on the spatial properties of vestibulospinal reflexes in decerebrate cats: role of the cerebellar anterior vermis. J Vestib Res 8:283–297.

Manzoni D, Pompeiano O, Bruschini L, Andre P (1999) Neck input modifies the reference frame for coding labyrinthine signals in the cerebellar vermis: a cellular analysis. Neuroscience 93:1095–1107.

Martin CZ, Brooks JX, Green AM (2018) Role of rostral fastigial neurons in encoding a body-centered representation of translation in three dimensions. J Neurosci 38:3584–3602.

Martin CZ, Lapierre P, Hache S, Lucien D, Green AM (2021) Vestibular contributions to online reach execution are processed via mechanisms with knowledge about limb biomechanics. Journal of neurophysiology 125:1022–1045.

Massion J (1994) Postural control system. Current opinion in neurobiology 4:877–887.

Maurer C, Mergner T, Peterka RJ (2006) Multisensory control of human upright stance. Experimental brain research 171:231–250.

Medendorp WP, Selen LJ (2017) Vestibular contributions to high-level sensorimotor functions. Neuropsychologia.

Meng H, May PJ, Dickman JD, Angelaki DE (2007) Vestibular signals in primate thalamus: properties and origins. J Neurosci 27:13590–13602.

Meng H, Blazquez PM, Dickman JD, Angelaki DE (2014) Diversity of vestibular nuclei neurons targeted by cerebellar nodulus inhibition. The Journal of physiology 592:171–188.

Merfeld DM, Zupan L, Peterka RJ (1999) Humans use internal models to estimate gravity and linear acceleration. Nature 398:615–618.

Merfeld DM, Young LR, Oman CM, Shelhamer MJ (1993) A multidimensional model of the effect of gravity on the spatial orientation of the monkey. J Vestib Res 3:141–161.

Mergner T, Rosemeier T (1998) Interaction of vestibular, somatosensory and visual signals for postural control and motion perception under terrestrial and microgravity conditions—a conceptual model. Brain Research Reviews 28:118–135.

Mildren RL, Cullen KE (2023) Vestibular contributions to primate neck postural muscle activity during natural motion. Journal of Neuroscience 43:2326–2337.

Moon IS, Kim JS, Choi KD, Kim M-J, Oh S-Y, Lee H, Lee H-S, Park S-H (2009) Isolated nodular infarction. Stroke 40:487–491.

Moreau-Debord I, Martin CZ, Landry M, Green AM (2014) Evidence for a reference frame transformation of vestibular signal contributions to voluntary reaching. Journal of neurophysiology 111:1903–1919.

Peterka RJ (2002) Sensorimotor integration in human postural control. Journal of neurophysiology 88:1097–1118.

Pouget A, Sejnowski TJ (1997) Spatial transformations in the parietal cortex using basis functions. J Cogn Neurosci 9:222–237.

Robinson DA (1963) A Method of Measuring Eye Movement Using a Scleral Search Coil in a Magnetic Field. IEEE transactions on bio-medical engineering 10:137–145.

Rochefort C, Lefort JM, Rondi-Reig L (2013) The cerebellum: a new key structure in the navigation system. Front Neural Circuits 7:35.

Rondi-Reig L, Paradis AL, Fallahnezhad M (2022) A Liaison Brought to Light: Cerebellum-Hippocampus, Partners for Spatial Cognition. Cerebellum (London, England) 21:826–837.

Ruggiero D, Batton RR, 3rd, Jayaraman A, Carpenter MB (1977) Brain stem afferents to the fastigial nucleus in the cat demonstrated by transport of horseradish peroxidase. The Journal of comparative neurology 172:189–209.

Salinas E, Abbott LF (1995) Transfer of coded information from sensory to motor networks. J Neurosci 15:6461–6474.

Salinas E, Abbott LF (1996) A model of multiplicative neural responses in parietal cortex. Proceedings of the National Academy of Sciences of the United States of America 93:11956–11961.

Shaikh AG, Meng H, Angelaki DE (2004) Multiple reference frames for motion in the primate cerebellum. J Neurosci 24:4491–4497.

Shaikh AG, Ghasia FF, Dickman JD, Angelaki DE (2005a) Properties of cerebellar fastigial neurons during translation, rotation, and eye movements. Journal of neurophysiology 93:853–863.

Shaikh AG, Green AM, Ghasia FF, Newlands SD, Dickman JD, Angelaki DE (2005b) Sensory convergence solves a motion ambiguity problem. Curr Biol 15:1657–1662.

Siebold C, Glonti L, Glasauer S, Buttner U (1997) Rostral fastigial nucleus activity in the alert monkey during three-dimensional passive head movements. Journal of neurophysiology 77:1432–1446.

Siebold C, Kleine JF, Glonti L, Tchelidze T, Buttner U (1999) Fastigial nucleus activity during different frequencies and orientations of vertical vestibular stimulation in the monkey. Journal of neurophysiology 82:34–41.

Siebold C, Anagnostou E, Glasauer S, Glonti L, Kleine JF, Tchelidze T, Buttner U (2001) Canal-otolith interaction in the fastigial nucleus of the alert monkey. Experimental brain research 136:169–178.

Smith MA, Majaj NJ, Movshon JA (2005) Dynamics of motion signaling by neurons in macaque area MT. Nature neuroscience 8:220–228.

Snyder LH, Grieve KL, Brotchie P, Andersen RA (1998) Separate body- and world-referenced representations of visual space in parietal cortex. Nature 394:887–891.

Stay TL, Laurens J, Sillitoe RV, Angelaki DE (2019) Genetically eliminating Purkinje neuron GABAergic neurotransmission increases their response gain to vestibular motion. Proceedings of the National Academy of Sciences 116:3245–3250.

Tarnutzer AA, Wichmann W, Straumann D, Bockisch CJ (2015) The cerebellar nodulus: perceptual and ocular processing of graviceptive input. Ann Neurol 77:343–347.

Walberg F, Dietrichs E (1988) The interconnection between the vestibular nuclei and the nodulus: a study of reciprocity. Brain research 449:47–53.

Wearne S, Raphan T, Cohen B (1998) Control of spatial orientation of the angular vestibuloocular reflex by the nodulus and uvula. Journal of neurophysiology 79:2690–2715.

Xing J, Andersen RA (2000) Models of the posterior parietal cortex which perform multimodal integration and represent space in several coordinate frames. Journal of cognitive neuroscience 12:601–614.

Yakusheva TA, Shaikh AG, Green AM, Blazquez PM, Dickman JD, Angelaki DE (2007) Purkinje cells in posterior cerebellar vermis encode motion in an inertial reference frame. Neuron 54:973–985.

Yoder RM, Taube JS (2014) The vestibular contribution to the head direction signal and navigation. Frontiers in integrative neuroscience 8:32.

Zhou W, Tang BF, King WM (2001) Responses of rostral fastigial neurons to linear acceleration in an alert monkey. Experimental brain research139:111–115.

Zipser D, Andersen RA (1988) A back-propagation programmed network that simulates response properties of a subset of posterior parietal neurons. Nature 331:679–684.

Zobeiri O, Cullen K (2022) Distinct representations of body and head motion are dynamically encoded by Purkinje cell populations in the macaque cerebellum. Elife 11:e75018–e75018.

Zupan LH, Merfeld DM, Darlot C (2002) Using sensory weighting to model the influence of canal, otolith and visual cues on spatial orientation and eye movements. Biological cybernetics 86:209–230.

